# Oncogenic *RAS* instructs morphological transformation of human epithelia via differential tissue mechanics

**DOI:** 10.1101/2021.01.19.427283

**Authors:** A. Nyga, J. Muñoz, S. Dercksen, G. Fornabaio, M. Uroz, X. Trepat, B. Baum, H. Matthews, V. Conte

**Affiliations:** Institute for Bioengineering of Catalonia (IBEC), The Barcelona Institute of Science and Technology (BIST), Barcelona – Spain; MRC Laboratory of Molecular Biology, Cambridge – United Kingdom; Department of Mathematics, Polytechnic University of Catalonia (UPC), Barcelona – Spain; Department of Biomedical Engineering, Eindhoven University of Technology (TU/e), Eindhoven – the Netherlands; Department of Physics, University of Barcelona (UB), Barcelona - Spain; Centro de Investigación Biomédica en Red en Bioingeniería, Biomateriales y Nanomedicina (CIBER-BBN), Barcelona – Spain; Department of Biomedicine, University of Barcelona (UB), Barcelona – Spain; Institució Catalana de Recerca i Estudis Avançats (ICREA), Barcelona – Spain; MRC Laboratory of Molecular Cell Biology, University College London (UCL), London – United Kingdom; Institute for Complex Molecular Systems (ICMS), Eindhoven University of Technology (TU/e), Eindhoven – the Netherlands

## Abstract

The RAS proto-oncogene is a critical regulator of cell state, morphology and mechanics, and plays a key role in cancer progression. Here, by using a human epithelial model in vitro, we ask how morpho-mechanical changes driven by oncogenic *RAS* activation at the level of individual cells are collectively integrated to drive changes in tissue behaviour. We found that the uniform oncogenic expression of *HRAS.V12* in confined epithelial monolayers causes reproducible changes in the structure and organization of the tissue, which acquires a transitory bilayered morphology. RAS-driven bilayering associates with reproducible layer-specific differences in cell-cell contractility and cell-matrix forces. These drive the initially flat tissues to form three-dimensional structures mimicking some of the behaviours seen in human cancers. Our findings establish a physical mechanism of cellular collectives through which uniform expression of RAS can be interpreted differently in different places of the same tissue to regulate its physiological and pathological morphology.

## Introduction

Epithelia are layered tissues, which provide separation between the inside (the stroma) and external milieu (the lumen). To perform this barrier function, the morphology of the epithelial tissues must be maintained as individual cells proliferate and die. Maintenance of epithelial homeostasis requires control of cell density through the dynamic regulation of cell proliferation, cell packing(*1*) and cell extrusion(*2*–*4*); the maintenance of a sheet-like morphology through the tight association of intercellular junctions(*5*); and stable anchoring between cells and the extracellular matrix(*6*). Intercellular junctions and matrix adhesions give epithelia the mechanical stability required to maintain the homeostatic layered architecture(*5, 6*). A critical role in the maintenance of epithelial homeostasis is played by the *RAS* proto-oncogene(*7, 8*).

The activity of the RAS family of GTPases, HRAS, NRAS and KRAS plays a critical role in the RAS-ERK signalling, which controls cell growth, division and survival. This pathway is dysregulated in a number of human diseases such as RASopathies(*9*). In addition, more than 50% of cancers involve the hyperactivation of the *RAS*/*RAF*/*MEK*/*ERK* pathway(*10, 11*), with more than 30% of all cancers being associated with specific activating mutations in *RAS* genes^8–10^. *RAS*-oncogenes disrupt epithelial homeostasis both in cell culture(*12*–*22*) (mostly studied by using clones) and in animal tissues(*17, 18, 23*–*27*). These studies show how *RAS*-transformed cells scattered within an epithelium may be segregated and expelled from the monolayer via extrusion or delamination(*12, 14, 19, 20, 22, 23*), through the mechanical engagement of the cellular interface with surrounding normal cells. Also, clusters of cells expressing oncogenic *RAS* or *SRC* (an upstream regulator of *RAS*(*28*)) can induce local dysplasia (*23, 24, 26, 27*), segregation(*16, 24*) or morphing(*23, 24, 26, 27*) of the affected area of the tissue as a consequence of differential mechanics along the extended interface between the non-transformed and transformed domains of the epithelial tissue.

These morpho-mechanical roles for *RAS* all rely on interfacial differences between juxtaposed non-transformed and *RAS*-transformed cells. Here, we have taken this further by monitoring, live, the impact of the uniform activation of oncogenic RAS on tissue morphology and mechanics. Our analysis reveals that uniform oncogenic RAS activation is sufficient to induce the separation of an epithelial monolayer into a multi-layered structure characterized by layer-specific differences in cell-cell contractility and cell-matrix adhesions. This transitory state of mechanical instability sets the tissue on a path to a 3D morphological transformation.

## Results

### Oncogenic *RAS-*expression induces the active dewetting of confined MCF10A monolayers

To systematically study the effects of *RAS*-activation on epithelial morphology and mechanics live (Fig. 1 A), we conditionally activated oncogenic *HRAS* in MCF10A human breast epithelial cells(*29*) (MCF10A/ER:HRAS^V12^). Rather than plating cells on stiff glass, for these experiments we used more-physiological(*30*) soft polyacrylamide-gel substrates with a stiffness of 12 kPa (see Materials & Methods) and a coating of rat tail collagen type-I micropatterned in circular shapes of 400 µm in diameter (Fig. 1 A). Epithelial cells were grown on these soft circular micropatterned substrates for 24 hours and either induced *HRAS*.*V12* expression by addition of 4-OHT (*RAS*-transformed) or DMSO for control (non-transformed). Surprisingly, we found that HRAS activation (Fig. 1B) was sufficient to transform the 2D epithelial monolayer into a 3D mass (Fig. 1 C-D and Fig. S1).

**Fig. 1.**
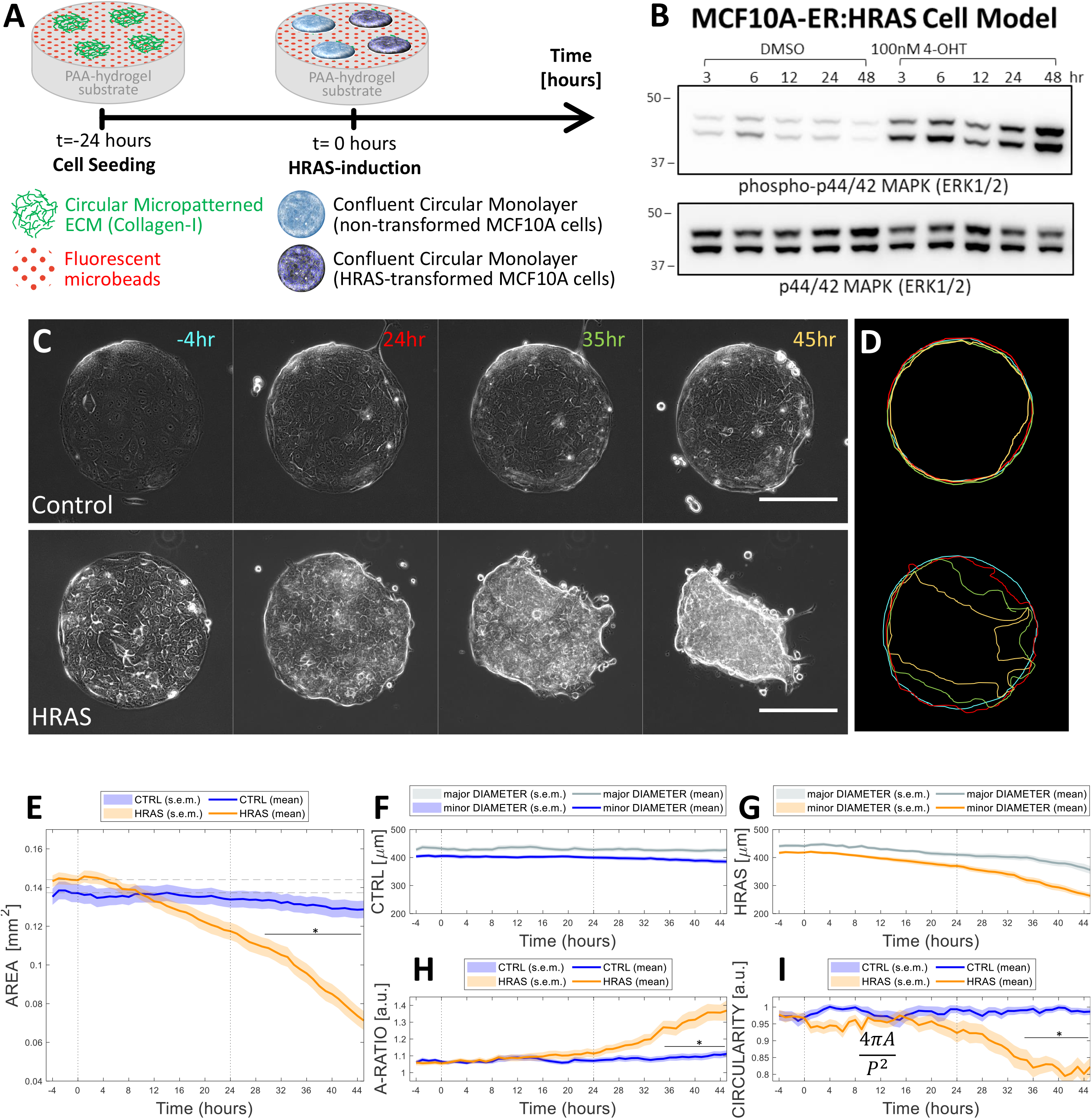
Morphological characterization of normal and *HRAS*-transformed MCF10A tissues. (**A**) Schematic of the experimental setup. (**B**) *HRAS* induction confirmed by increase in phosphorylation of MAPK (ERK1/2) shown in representative Western Blots. (**C**) Phase contrast time-lapse of selected non-transformed (control) and *HRAS*-transformed MCF10A monolayers (imaging starts at t=-4 hours and *HRAS*-activation is induced at t=0 hours; scale bar is 200 µm). (**D**) Contours of epithelia shown in panel C (time progresses with blue, red, green and yellow colours). (**E**) Time-evolution of the surface area of the epithelia’s footprint on the substate matrix (epithelial domain); (**F-G**) time-evolution of the major and minor diameters of the epithelia domain in the case of (**F**) non-transformed and (**G**) HRAS-transformed tissues – these quantities are defined as the axes of the elliptical envelope having the same normalized second central moments as the epithelial domain; (**H**) time-evolution of the epithelial domain’s aspect ratio – this quantity is defined as the major axis divided by the minor axis of the epithelial domain’s (as defined in subpanels F-G); (**I**) time-evolution of the epithelial domain’s circularity – this quantity is defined as 4π times the area (as defined in subpanel E) divided by the squared perimeter of the epithelium’s domain. (**E-I**) Statistics over 15 non-transformed epithelia and 16 HRAS-transformed epithelia from at least 4 independent experiment repeats. Each epithelium was imaged for at least 50 hours. Median over each epithelial domain at each time point of its evolution. Time-evolution graphs represent Mean±S.E.M. of medians at each time point. 2-way ANOVA with Bonferroni post-test, *p<0.05.

To define the path of this 2D-to-3D tissue transformation, we monitored the process via a set of objective morphological (Fig. 1) and mechanical measures (Fig. 2). Specifically, we quantified the area (Fig. 1 E), aspect ratio (Fig. 1 F-H), circularity (Fig. 1 I) and traction forces (Fig. 2) for each cellular island.

**Fig. 2.**
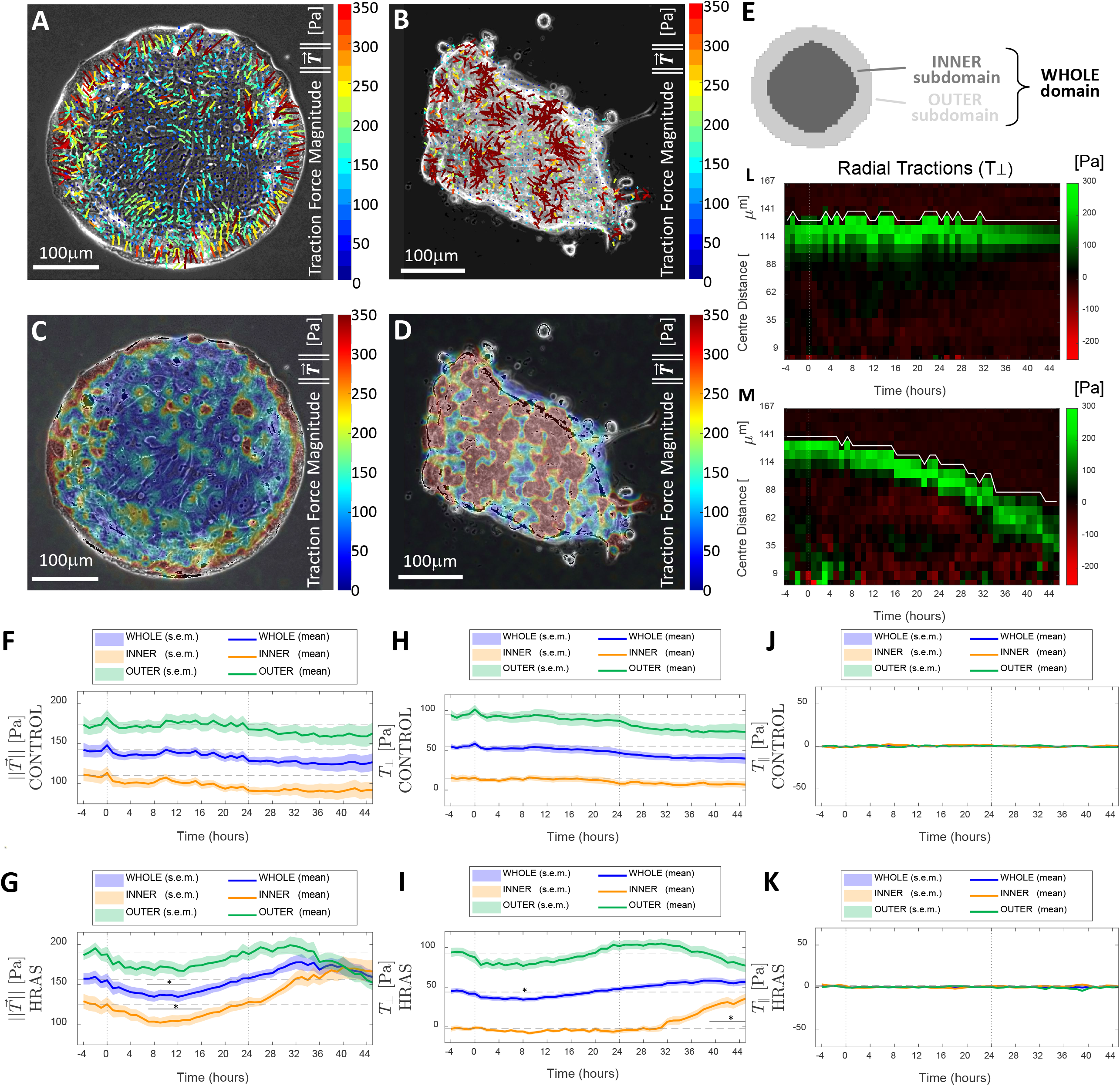
Mechanical characterization of normal and *HRAS*-transformed MCF10A tissues. Overlays of traction-force vectors (**A-B**) and traction-force magnitude maps (**C-D**) on phase contrast images of confined MCF10A epithelia; (**A**,**C**) representative non-transformed epithelium at t=19 hours; (**B**,**D**) representative *HRAS*-transformed epithelium at t=45 hours. (**E**) Schematic representing the whole epithelial domain along with its outer and inner subdomains. (**F-K**) Time evolution of magnitudes and components of the traction-force field in the whole epithelial domain (blue) as well as in the inner (orange) and outer (green) epithelial subdomains. (**F-G**) Time evolution of the average Traction-field ±magnitude for (**F**) non-transformed epithelia and (**G**) *HRAS*-transformed epithelia. (**H-K**) Time evolution of the average traction force components: (**H-I**) perpendicular to the epithelial domain’s edge (T_⊥_) for (**H**) non-transformed and (**I**) *HRAS*-transformed epithelia; (**J-K**) tangential to the epithelial domain’s edge (T_‖_) for (**J**) non-transformed and (**K**) *HRAS*-transformed epithelia. (**L-M**) Kymographs of the radial (perpendicular) component of the traction field T_⊥_: (**L**) for a representative non-transformed MCF10A epithelium and (**M**) for a representative *HRAS*-transformed MCF10A epithelium. White lines represent the average evolution of the edge of the island in time, the centre of the island colocalizing with the bottom of the graph. A negative component T_⊥_ (red) means that the corresponding traction-force vector is in oriented towards the exterior of the epithelial domain’s edge, whereas a positive component T_⊥_ (green) is indicative of traction-force orientation toward the interior of the epithelial domain’s edge. (**F-K**) Statistics over 15 non-transformed epithelia and 16 *HRAS*-transformed epithelia from at least 4 independent experiment repeats. Each epithelium was imaged for at least 50 hours. Median over each epithelial domain at each time point of its evolution. Time-evolution graphs represent Mean±S.E.M of medians at each time point. 2-way ANOVA with Bonferroni post-test, *p<0.05.

Morphological analyses revealed differences between RAS and control tissues beginning ∼4 hours after *HRAS* induction. Whereas control tissues remained flat and maintained a near constant area throughout the time window of our analysis, from 4 hours after oncogenic *HRAS* induction, tissue islands exhibited a slow decrease in area. This decrease in circular area continued in an approximately symmetric fashion up until 24 hours after *HRAS* induction (Fig. 1 F-H), after which the symmetry of *RAS*-transformed islands started to break (Fig. 1 F-H). At this point, *RAS*-transformed tissues underwent a loss of tissue circularity and an increase in aspect ratio (Fig. 1 H-I). This was followed by a rapid transition as the 2D *RAS-*transformed epithelium started to morph into a 3D tissue (Fig. S1). None of these changes were seen in control tissues.

Mechanical analyses carried out using Traction Force Microscopy(*31*) showed that traction forces of higher magnitude were mainly localized at the periphery of both non-transformed and *RAS*-transformed tissues (Fig. 2 A, F-G). In both cases, the elastic strain energy actively transferred by the epithelium to the underlying substrate also followed a similar trend (Fig. S2). The concentration of traction force at the periphery of circular epithelia is in line with previous theoretical(*32, 33*) and experimental studies carried out on epithelial colonies of human colon carcinoma (HCT-8)(*34*) and on confined circular monolayers of canine kidney cells (MDCK)(*35, 36*).

We noticed that high traction forces tended to be more ordered at the epithelium’s periphery (Fig. 2 A,B). Thus, we turned our attention to the topology of the traction field by further quantifying the distribution and orientation of traction forces within the different quadrants of the epithelial domain’s (Supplementary Text 1). Our analysis showed that high traction forces at the epithelial domain’s periphery appeared to be oriented towards the epithelial domain’s centre, whereas lower traction forces underneath the bulk of epithelia were less organised in both non-transformed and *HRAS*-transformed tissues (Fig. S3). In order to decompose the traction field along the directions normal and tangential to the epithelial domain’s edge (Fig. S4), we computed: i) the net radial (T_⊥_) and net tangential (T_‖_) traction force components as a function of time in the whole epithelial domain as well as in the central (inner) and peripheral (outer) domains (Fig. 2 E). These time-trends (Fig. 2 H-K) confirmed that the net physical interactions at the interface between epithelia and substrate developed along the direction perpendicular to the epithelium’s edge for both non-transformed and *HRAS*-transformed epithelia throughout the analysis.

To further average out spatial-temporal fluctuations of the traction force field and to better visualize reproducible force patterns, we averaged the net radial traction components T_⊥_ in space, along lines concentric to the edge of the island. Upon displaying these space-averages as a function of distance from the epithelial domain’s edge, we obtained kymographs of the net radial traction component T_⊥_ for both non-transformed (Fig. 2 L) and *RAS*-transformed MCF10A epithelia (Fig. 2 M). Kymographs confirm that it is the intense net radial-traction components that concentrate mainly at the periphery of both non-transformed and *RAS*-transformed epithelia throughout the experiment. By analysing the stress distribution within the epithelial domain through Monolayer Stress Microscopy(*31*) (Materials&Methods), we further showed that non-transformed tissues successfully establish and maintain epithelial homeostasis by means of long-range transmission of physical forces throughout the monolayer from opposite locations of the epithelial domain’s edge, which results in tension accumulating throughout the bulk of the epithelial domain (Fig. S5).

Our traction force analysis also showed that, unlike control epithelia, the average intensity of the traction field of *RAS*-transformed epithelia undergoes a characteristic two-phase oscillation in the time window of our analysis (Fig. 2 G). This was observed as a decreasing-increasing phase over the first 24 hours following *HRAS* induction – when average traction forces dropped for approximately 8 hours before recovering again by approximately 24 hours (Fig. 2 G-I and Fig. S3). This disrupts the mechanical homeostasis of intact non-transformed circular epithelia and sets them on course of an abrupt symmetric decrease in epithelial area within the first 24 hours of oncogene activation (Fig. 1 E) followed by a 2D-to-3D morphological transformation (Fig. 1 B and Fig. S1).

As expected, the overexpression of *KRAS*^V12^ in confined circular epithelia respectively led to similar results (Fig. S6-S7). Upon *KRAS*^V12^-activation, confined circular monolayers of inducible MCF10A/ER:KRAS^V12^ cells followed a similar morpho-mechanical fate to those of MCF10A/ER:HRAS^V12^ cells, suggesting that different RAS isoforms *HRAS* and *KRAS* have similar morpho-mechanical effects (Fig. 2 and Fig. S3,S6-S7).

The relatively rapid morphological (Fig. 1) and mechanical (Fig. 2) changes observed in confined transformed MCF10A monolayers from t=24 hours onward are characteristic of a cellular process known as active dewetting. In a previous case(*37*), this has been associated with a monotonic increase in centripetal traction forces linked to the retraction of a circular confined monolayers of human-breast adenocarcinoma cells (MDA-MB-231). This led us to hypothesize that oncogenic *RAS* expression primed tissues for dewetting during the first 24 hours of *RAS* expression by inducing structural and mechanical changes within the monolayer while still morphologically flat. To test this, we turned our focus to the changes in tissue organization during the first 24 hours of cellular evolution from *RAS-*activation.

### Oncogenic *HRAS* expression triggers the bilayering of confined MCF10A monolayers

By analysing movement of the tissue in the z-plane orthogonal to the underlying substrate, we observed early changes in the morphology of transformed tissues that were evident in measures of both nuclear and monolayer height (Fig. 3 A-B). Thus, within 24 hours of oncogene activation RAS-transformed tissues became an average of 25% thicker than non-transformed ones (Fig. 3 A and Fig. S8). This was accompanied by an increase in cell packing that was markedly higher in the confined *RAS-*transformed monolayers, leading to the observed decrease in the area of individual cells (Fig. 3 C). Both non-transformed and *RAS-*transformed MCF10A monolayers presented uniform nuclear heights up to 8 hours from oncogene activation (Fig. 3 B). Thereafter and up until 24 hours, the nuclear heights of *RAS-*transformed tissues showed greater heterogeneity, an indicator of the presence of topological heterogeneities within the confined epithelium(*38*). Indeed, the RAS-expressing epithelium (but not the control) became multi-layered over this time period (Fig. 3 D). Confocal microscopy revealed how confined *RAS-*transformed epithelia segregated into two discrete tissue layers with very distinct organizations (Fig. 3 D-E). The cells forming the top layer of *RAS-*transformed tissues were flatter and more spread than those in the bottom layer (Fig. 3 E), having undergone a 1.5-fold increase in cell perimeter (Fig. 3 G) and a 3-fold average increase in the cell area (Fig. 3 F) without any substantial effect on cell shape – the cell aspect ratio remained approximately constant in both layers (Fig. 3 H). We next turned our focus on the cellular alterations that could cause the bilayering.

**Figure 3.**
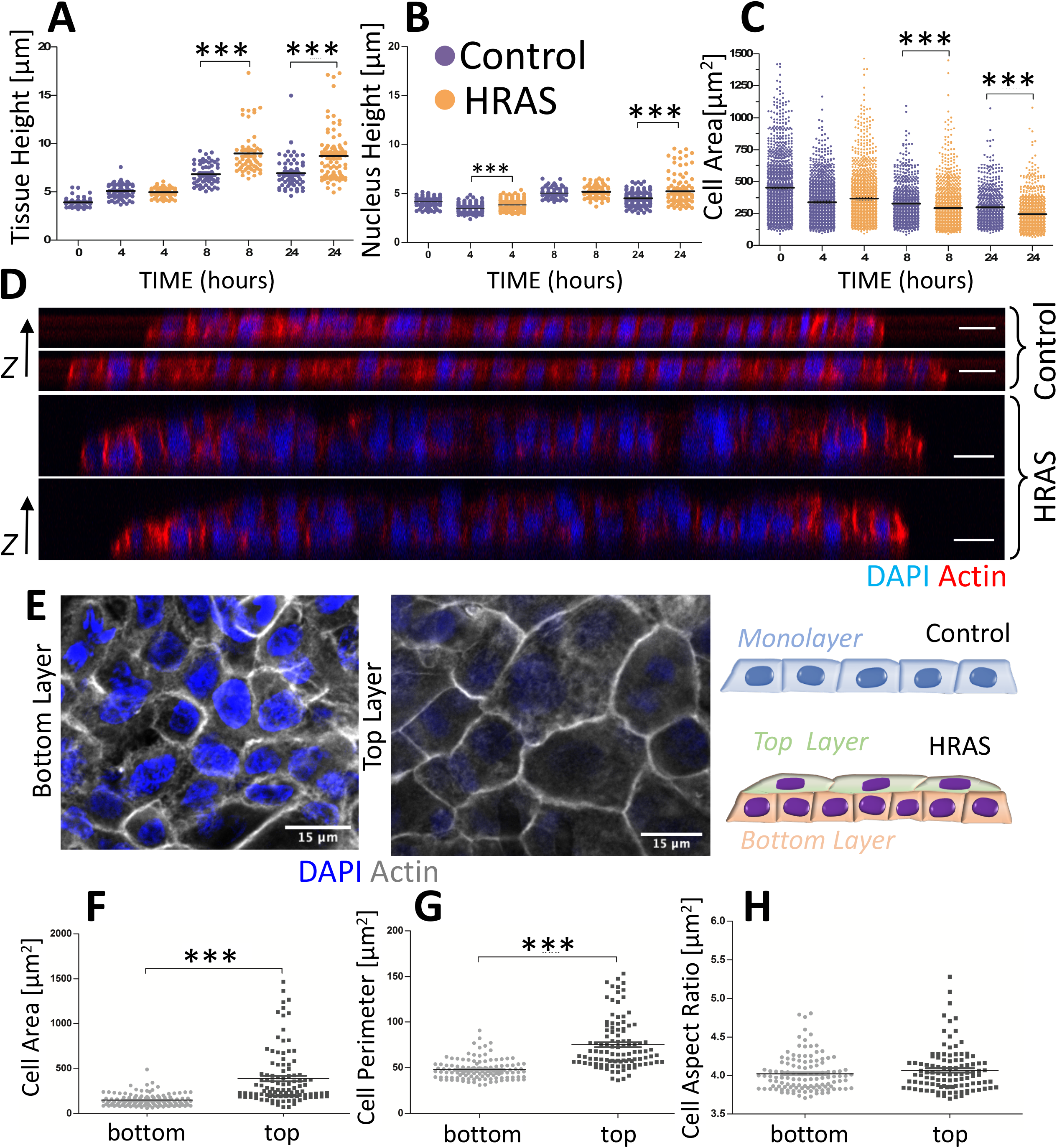
Oncogenic *HRAS*-expression induces the bilayering of MCF10A monolayers. (**A-C**) Measurements of epithelial monolayers features at selected time points for both non-transformed and *HRAS-*transformed epithelia: (**A**) tissue height, (**B**) nucleus height and (**C**) cell area (mm^2^). Kruskal-Wallis statistic test with Dunn’s Multiple Comparison Test. (**D**) Representative confocal images of non-transformed monolayer (top-two subpanel) and *HRAS-*transformed bilayer (bottom-two subpanel) at t=24 hours from oncogene induction – focal planes is orthogonal to the substrate matrix (z-axis). Actin is labelled in red and DAPI in blue. Scale bar = 20 µm. (**E**) Representative confocal image of non-transformed monolayer and *HRAS*-transformed bilayer – focal plane crosses the tissue parallelly to the substrate matrix. Actin is labelled in shades of grey and DAPI in blue. Scale bar = 15 µm. (**F-H**) Quantification of cell features in the top and bottom layers of *HRAS*-transformed bilayers: (**F**) cell surface area, (**G**) cell surface perimeter and (**H**) cell shape index. Mean±S.E.M. of median values from at least 3 individual patterns. Mann Whitney test, *** p<0.001.

### Oncogenic *RAS*-expression induces layer-specific differences in cell-cell contractility and cell-matrix adhesions of the confined MCF10A bilayers

Stable RAS transformation has been shown to disrupt cadherins to promote cell invasion(*39*–*41*). Therefore, we hypothesized that the bilayering induced by oncogenic RAS-expression might also be caused by disruption to cell-cell junctions. Non-transformed MCF10A cells constitutively expressed E-cadherin and, thus, exhibited uniform cell-cell junctions throughout the tissue. Strikingly, E-cadherin expression did not change during *RAS-*activation (Fig. 4 A). Nevertheless, we observed differences in the localization of E-cadherin in the two layers of the developing bilayer (Fig. 4 B). Cells in the bottom layer tended to have relatively low levels of E-Cadherin at cell-cell junctions, while junctional E-cadherin levels remained similar to those before *RAS* activation in the top layer (Fig. 4 C). In addition, the ratio of junctional to cytoplasmic E-cadherin was decreased in the bottom layer (Fig. 4 D), suggestive of a redistribution of the protein from junctions to cytoplasm. Intriguingly, this change in E-cadherin localization in the bottom layer induced by the *RAS-*transformation was paralleled by a reduction of cell-matrix adhesion between the epithelial cells of this layer and the hydrogel substrate, as shown by a decrease in the expression of the collagen receptor integrin β1 (Fig. 4 E-G).

**Figure 4.**
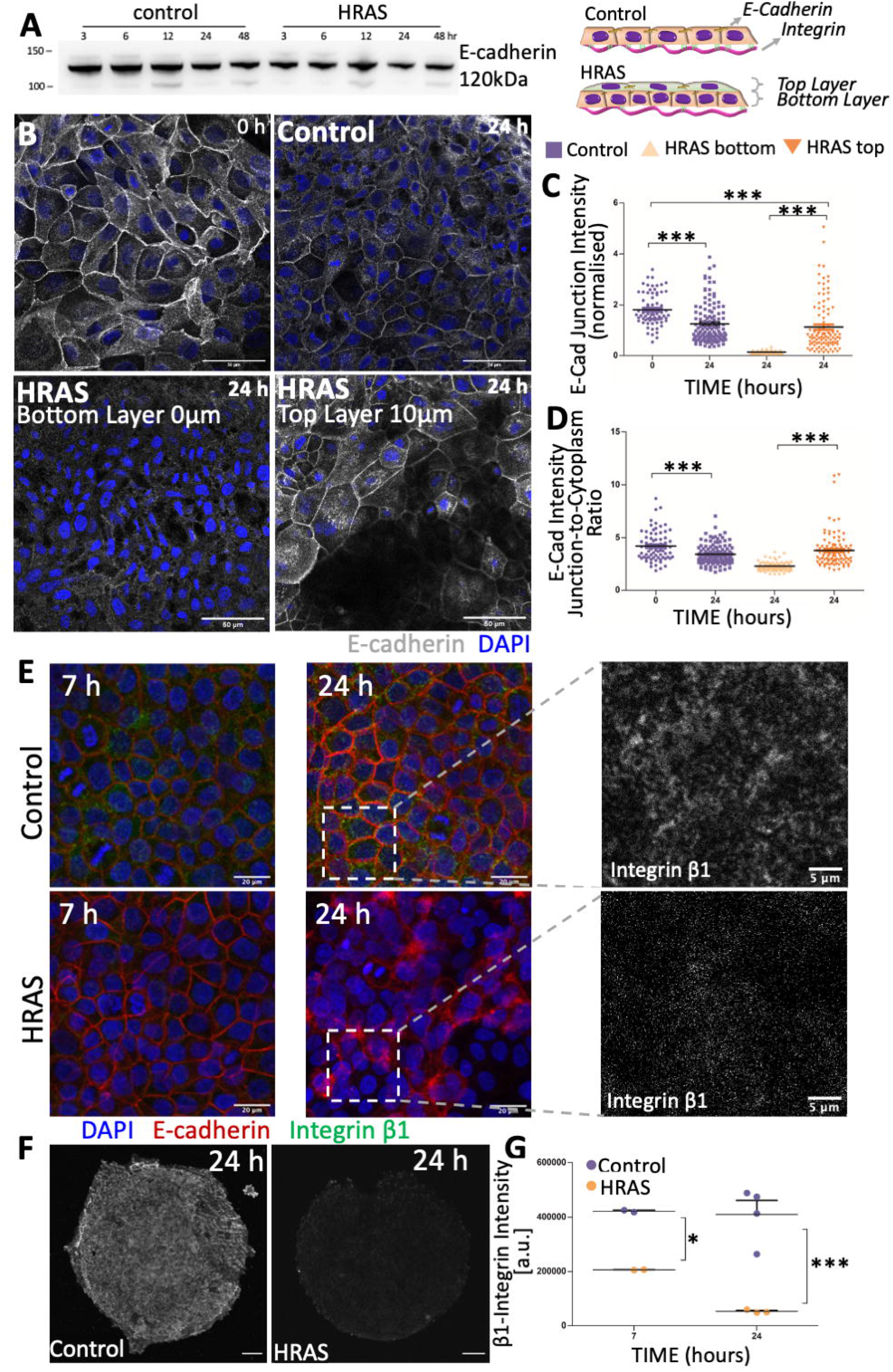
Oncogenic HRAS-expression alters the expression of E-cadherin and ß1-integrin. (**A**) Western Blot showing E-cadherin expression in non-transformed (control) and *HRAS*-transformed MCF10A cells over 48 hours from oncogene induction. (**B**) Representative confocal images showing the distribution of E-cadherin at t=0 hours and within non-transformed and *HRAS*-transformed tissues at t=24 hours. E-cadherin labelled in grey, DAPI in blue. Scale bar = 50 µm. (**C-D**) Global E-cadherin fluorescence in non-transformed (control) and *HRAS*-transformed epithelia: (**C**) junctional intensity (normalized) and (**D**) Ratio of junctional to cytoplasmic intensity. (**E**) Confocal images of non-transformed and *HRAS*-transformed epithelia after 7 and 24 hours of oncogene induction stained for E-cadherin (red), DAPI (blue) and integrin ß1 (green in panels at the centre and shades of greys in panels on the right). Scale bar = 20 µm (**F**) Confocal images of entire non-transformed monolayer and *HRAS*-transformed bilayer after 24 hours of oncogene induction stained for integrin ß1 (shades of greys) – focal plane crosses the tissue parallelly to the substrate matrix. Scale bar = 50 µm. (**G**) Intensity of global ß1-integrin fluorescence in non-transformed and *HRAS*-transformed epithelia after 7 and 24 hours of oncogene induction. **E-cadherin:** Mean±S.E.M. of median values from at least 3 individual patterns and Kruskal-Wallis statistic test with Dunn’s Multiple Comparison Test. **Integrin:** Mean±S.E.M.; 2-way ANOVA with Bonferroni post-test. *p<0.05, **p<0.01, ***p<0.001.

The structural alterations occurring in *RAS*-transformed epithelia at the level of cell-cell junctions (Fig. 4 A-E) and cell-matrix adhesions (Fig. 4 E-G) correlated with the initial decrease observed in traction forces transferred by the tissue to the substrate matrix during the first ∼8 hours of oncogene induction (Fig. 2 G). However, traction forces recovered to pre-transformation levels by 24 hours of *RAS*-transformation (Fig. 2 G), while adhesion to the substrate kept decreasing without recovering (Fig. 4 E-G). Therefore, we hypothesized that active tension within the tissue might also be affected by the *RAS*-transformation. To further understand whether *RAS*-driven alterations to tissue structure also reflected on tissue tension (driven by the contractile cellular actomyosin cortex(*42, 43*)), we studied the distribution of key cortex components F-actin and pMLC2(*44*) in both non-transformed and *RAS-*transformed tissues. While non-transformed epithelia show uniform distribution of F-actin and pMLC2 (Fig. 5 A-C), oncogenic *RAS* expression induced a gradient in the distribution of F-actin and pMLC2 throughout the bilayer within 24 hours of oncogene activation (Fig. 5 D-H). F-actin and pMLC2 fluorescent intensity were increased at the periphery of the bottom layer of *RAS-*transformed bilayers (Fig. 5 D-F). This fluorescent intensity was lower and more heterogeneously distributed throughout the top layer (Fig. 5 G-H), although spots of highly increased pMLC2 expression were visible at the periphery and in the middle of top layer of *RAS-*transformed bilayers (Fig. 5 G). Unexpectedly, these results show that while non-transformed monolayers retain a uniform homogenous state of tension (Fig. S5) and organization (Fig. 1-2) as they grow under these conditions, the uniform expression of oncogenic *RAS* for 24 hours leads to the establishment of a tension gradient that destabilizes the tissue. Thus, *RAS-*transformed circular epithelia underwent a significant radially symmetric contraction (Fig. 1, E-I) and formed multiple layers with very different properties (Fig. 2). We hypothesised that these *RAS*-induced alterations to cell-cell junctions, cell-matrix junctions and tissue tension prime tissues for active dewetting. To test whether this is likely to be the case, we developed a simple computational model of circular cellular tissues.

**Figure 5.**
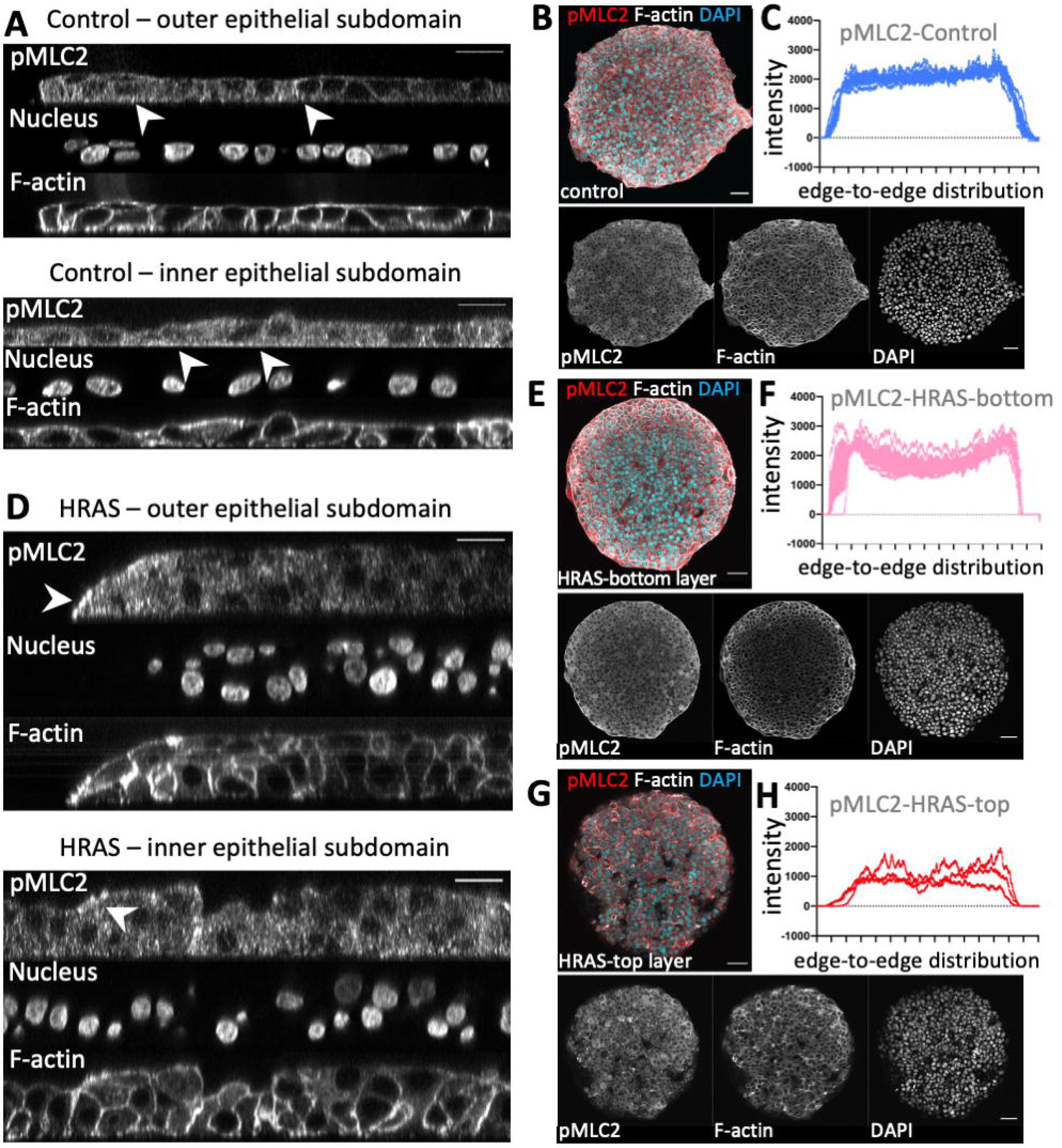
Oncogenic *HRAS*-induction causes a tension differential in the MCF10A bilayers. Representative confocal images of non-transformed monolayer (**A**) and *HRAS-*transformed bilayer **(D)** at t=24 hours from oncogene induction stained for pMLC2, F-actin and DAPI (shades of greys) – focal planes are orthogonal to the substrate matrix. Scale bar = 20 µm. (**B**,**E**,**G**) Representative confocal images of entire non-transformed monolayer (**B**) and *HRAS*-transformed bilayer (**E**, bottom layer and **G** top layer of the bilayer) – focal plane crosses the tissue parallelly to the substrate matrix. pMLC2, F-actin and DAPI (nuclei) are labelled either in colour code (red, grey and blue respectively) or shades of greys. Scale bar = 50 µm (**C**) Averaged global intensity of pMLC2 fluorescence from edge to edge of the circular epithelial domain of non-transformed monolayers (n=3 and mean±S.D. in blue) (**F**,**H**) Averaged global intensity of pMLC2 fluorescence from edge to edge of the circular epithelial domain of HRAS-transformed bilayers (n=3): (**F**) mean±S.D. of bottom layers (in pink); and (**H**) individual intensity profiles for the three top layers (in red).

### Oncogenic *RAS*-expression makes confined monolayers mechanically instable

The circular tissue bilayers were modelled as a 2D continuum elastic disk with finite thickness in two-dimensional plain-stress approximation (Fig. 6 A, Materials & Methods). The tissue *in silico* is mechanically coupled to the substrate matrix at discrete focal contact points (Fig. 6 B). Elastic friction laws were used to emulate the dynamics of discrete focal contact points between cells of the monolayer and the substrate matrix in the simplest possible way(*45*) (Fig. 6 B). The finite element method was used to resolve tissue motion generated against friction with the substrate by active contractile cellular forces (Fig. 6 C-D and Materials & Methods). We assumed that local cellular contractility within the tissue was proportional to the relative fluorescence intensities of pMLC2 (Fig. 5 – Materials & Methods). The effects of *RAS-*induction within tissues were modelled based on experimental data as a local drop in cell-matrix adhesion from baseline pre-transformation levels (Fig. 6E – based on data in Fig. 4 G); and, concomitantly, a local rise in tissue tension (cell contractility) from baseline pre-transformation levels (Fig. 6 F – as observed in Fig. 5 C,F,H).

**Figure 6.**
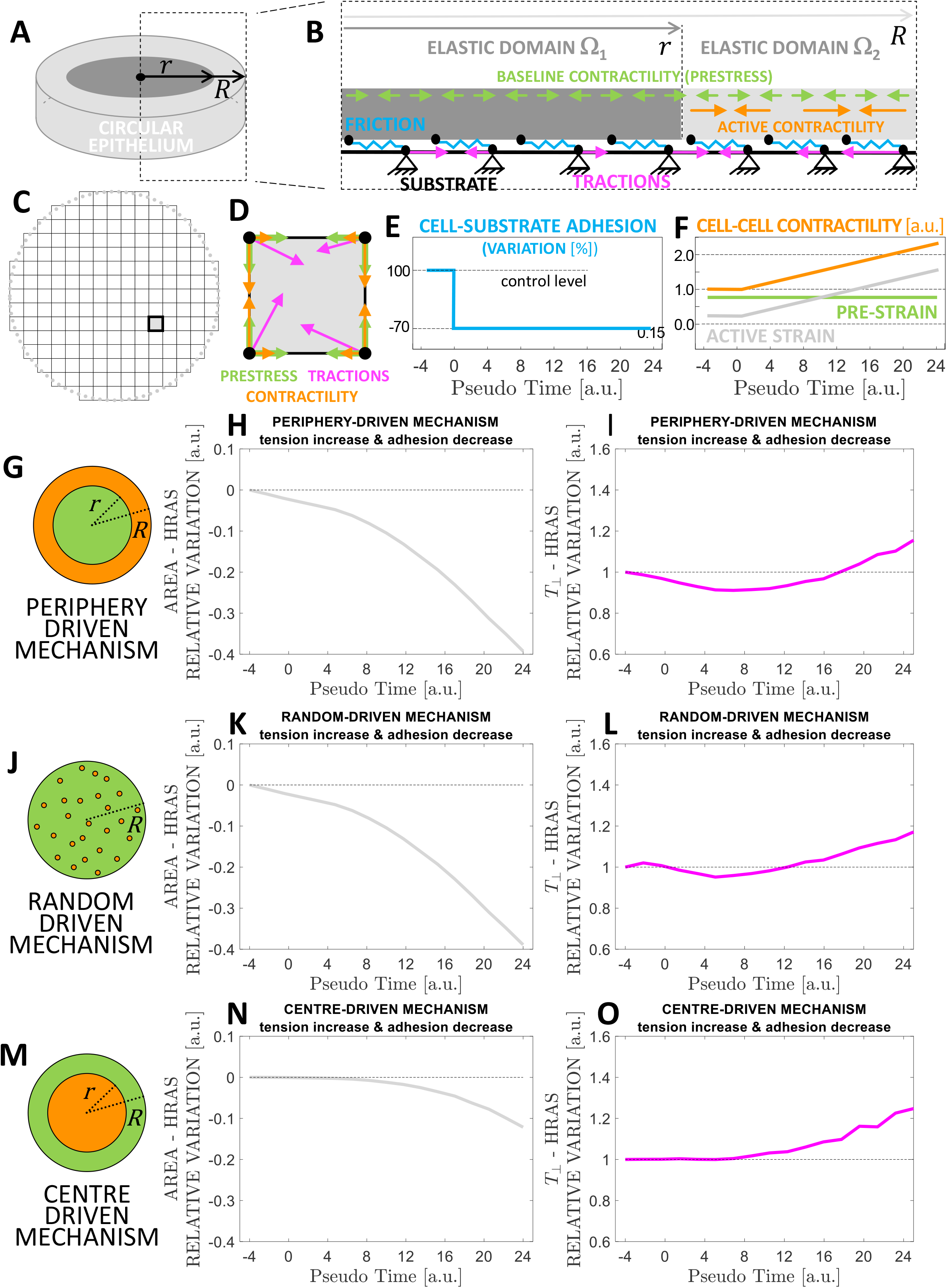
*In silico* model of *HRAS*-driven mechanical instability in MCF10A circular tissues. (**A-B**) Schematic representing the experimental features captured and emulated by the computational model. (**A**) The epithelium is represented as a 2D continuum (bulk stiffness *E* and Poisson Ratio *v*) having finite thickness in plane-stress approximation (grey). (**B**) Schematic representing the inter- and intra-cellular forces along with the cell-matrix forces at the interface with the elastic substrate. The entire domain of the epithelium is uniformly subjected to a contractile pre-strain constant in time (green arrows, Materials&Methods). A subdomain Ω_2_ of the epithelial domain (light grey) can also develop further contractile strain (active tension) – the distribution of tension within the epithelium represented here corresponds to that one illustrated in panel G. Active tensions and pre-strain equilibrate passive elastic forces within the epithelium and result in traction forces (magenta arrows) being transferred to the substrate matrix via elastic friction (blue springs). (**C**) Finite element discretization of the 2D epithelium with a representative finite element represented in dark grey and outlined in black. (**D**) Forces acting at nodes of a representative finite element. (**E**) Adhesion with substrate stays at pre-transformation levels in correspondence with the epithelial subdomain Ω_1_ that is subject to pre-strain only whereas it decreases by 70% in correspondence with the epithelial subdomain Ω_2_, which is subject to additional active tension. (**F**) Active intercellular tension increases monotonically within the epithelial subdomain Ω_2_. (**G**,**J**,**M**) Schematics illustrating the different topologies according to which active tension can locally increase within the tissue (subdomains Ω_1_ and Ω_2_ are color-coded in green and orange respectively). (**H**,**K**,**N**) Epithelial surface area trends in correspondence to each of the scenarios in panels G-I respectively. (**I**,**L**,**O**) Epithelial traction force trends in correspondence to each of the scenarios in panels G-I respectively.

We considered 3 different topological scenarios to locally configure tension and adhesion. These were: i) a periphery-driven mechanism (Fig. 6 G-I), whereby increased cell-cell tension (Fig. 6 F) and decreased cell-matrix adhesion (Fig. 6 E) arise from cell contractility at the periphery of the circular epithelium (Fig. 6 G); ii) a random mechanism (Fig. 6 J-L), whereby increased cell-cell tension (Fig. 6 F) and decreased cell-matrix adhesion (Fig. 6 E) arise from cell contractility at randomly scattered locations throughout the epithelium(Fig. 6 J); and iii) a centre-driven mechanism (Fig. 6 M-O), whereby increased cell-cell tension (Fig. 6 F) and decreased cell-matrix adhesion (Fig. 6 E) arise from cell contractility at the centre of the circular epithelium (Fig. 6 M). We used finite element methods to model the mechanics of the tissue under the respective conditions (Fig. 6 C-D, Materials & Methods and Supplementary Text 2). We then computed the profiles of the area of the circular tissue (Fig. 6 J-L) and the traction force transmitted by the tissue to the substrate (Fig. 6 M-O) in all three topological scenarios detailed above as a function of cellular contractility and/or cell-matrix adhesion (Fig. 6 G-O and Fig. S9-S10). In all three topological scenarios (Fig. 6 G-I), simulations confirmed that: i) local increases in cellular contractility (and, thus, global increases in tissue tension) are sufficient to explain the symmetric contraction of tissue area and the increase in traction-force intensity (Fig. S10); ii) the reduction in cell-matrix adhesion alone is sufficient to account for the reduction in traction-force intensity (Fig. S9); iii) the combination of the decrease in cell-matrix adhesion with the increase in active tissue tension can result in an oscillation of the traction force intensity (Fig. 6 I) of the kind we observed experimentally (Fig. 2 G,I) in conjunction with symmetric tissue retraction (Fig. 6 H and Fig. 1 E, H, I). This oscillation could not be produced by the centre-driven mechanism and was particularly pronounced in the periphery-driven mechanism (Fig. 6 G) – the latter more closely reflecting the experimental distribution of active contractility (F-actin and pMLC2) in *RAS*-transformed bilayers (Fig. 5 E-H).

Taken together, our computational analyses showed that the combination of the reduction in adhesion and the ensuing redistribution of active tension within *RAS-*transformed tissues towards their periphery is sufficient to induce a state of mechanical instability in these tissues during the first 24 hours of oncogene induction. This instability sets the transformed tissues on a path of symmetric contraction which effectively primes them for active dewetting from that point onward (Fig. 1).

## Discussion

The *RAS*-genes superfamily functions as a crucial signalling hub within the cell, which controls several of its critical signalling cascades. This also reflects how profoundly the somatic and germline abnormal activity of *RAS* affects human development and disease(*9*–*11*). *RAS* mutation alters the morphology and mechanics of individual cells(*46*), and here we showed how these morpho-mechanical changes can collectively integrate at the tissue scale to instruct the abnormal morphing of epithelia (Fig. 1). Here, we have used a combined experimental and computational analysis to ask how *RAS*-driven changes in cellular morphologies and forces affect the homeostasis of intact confined MCF10A epithelial monolayers (Fig. 1,2). Our analysis shows, quite surprisingly, that uniform *RAS*-induction can lead to the formation and contraction of a complex 3D tissue architecture that undergoes active dewetting (Fig. S1).

This morpho-mechanical transformation is mediated by a series of changes in epithelial architecture and mechanics that are induced during the first 24 hours of uniform *RAS-*oncogene induction (Fig. 3). These include: i) the establishment of two layers of cells that have very different patterns of a E-cadherin localization, with high E-cadherin cell-cell junctions in the top layer and low E-cadherin cell-cell junctions in the bottom layer (Fig. 4); ii) local reductions in cellular adhesion to the substrate (Fig. 4), leading to decreased traction-forces transferred by the affected tissues to the substrate (Fig. 2); and, iii) a redistribution of regulators of cellular tension (F-actin and pMLC2), which become concentrated at the periphery of the bilayer (Fig. 5). In the future it will be fascinating to assess how these local changes in cell biology are induced by the uniform expression of oncogenic *RAS*. Our analysis shows how these local heterogeneities, when combined, give rise to the change in overall tissue organization and mechanics.

Increased intercellular tension has been previously reported as a key driver for the active dewetting of circular monolayers of human breast adenocarcinoma cells (MDA-MB-231) *in vitro*(*37*). In this previous study, tension differential was induced via the upregulation of E-cadherin – which is not normally expressed by transformed MDA-MB-231 cells, unlike our MCF10A cells. Within approximately 25 hours of an increase in E-cadherin expression in MDA-MB-231 monolayers, these epithelia underwent a dramatic increase of intra-and inter-cellular tension that triggered the decrease in tissue area and subsequent 2D-to-3D morphological transition of dewetting(*37*). Importantly, changes in E-cadherin levels alone could not account for tissues’ active dewetting. Instead, a decrease in cell-matrix adhesion to the underlying substrate affected the cell-cell tension threshold that regulated the temporal onset of active dewetting of an island with a given initial diameter(*37*).

While the dewetting phenomenon that we observe is similar, the cause is very different in our system. In the case of circular MCF10A monolayers, the homeostatic balance between cell-matrix adhesion (Fig. 2) and cell-cell tension (Fig. S5) allows intact non-transformed monolayers to preserve their morphology throughout the time window of our analysis (Fig. 1). However, the uniform expression of oncogenic *RAS* disrupts this homeostatic equilibrium by eliciting non-uniform architectural (Fig. 3), structural (Fig. 4) and mechanical changes (Fig. 5) across the tissue. Our simulations show that the imbalance in cell-cell tension and cell-matrix adhesion brought about by oncogenic *RAS-*expression are sufficient to place transformed tissues into a state of mechanical instability (Fig. 6), which primes the tissues for active dewetting (Fig. 1). Mechanical imbalance following the oncogenic expression of *RAS* has been previously shown to drive the pathological morphing of the pancreatic duct(*26*). There, simulation and experiment showed that oncogenic *RAS*-expression affected the mechanical balance between the non-transformed and the *RAS*-transformed domains of the duct by levelling the tensional gradient between the apical and basal sides of the *RAS*-transformed subdomain of the duct(*26*). Here, uniform oncogenic *RAS*-expression throughout the tissue leads to tensional imbalance and disruption of homeostatic equilibrium by inducing the formation of two tissue layers with very different adhesive and contractile properties. Importantly, by establishing a mechanical imbalance through *RAS*-transformed tissues, epithelial bilayering may also provide a favourable intermediate mechanism to abnormally morph tissues without the need for an extended interface with non-transformed tissues(*16, 23, 24, 26, 27*). Overall, our findings establish a new physical mechanism of cellular collectives through which *RAS* can regulate the physiological and pathological morphology of epithelia in human development and disease(*9*–*11*) autonomously of cell competition.

## Materials and Methods

### MCF10A cell culture

Immortalized epithelial breast cell line MCF10A was transfected with inducible ^12^V-mutated form of the *HRAS* gene (a gift from Julian Downwards lab UCL, London, UK)(*29*), referred to as MCF10A/ER.HRAS V12. They were maintained in complete medium composed of: phenol-free DMEM-F12 medium (ThermoFisher, #11039047) supplemented with 5% charcoal-stripped horse serum (ThermoFisher, #16050122), 100 U/ml penicillin and 100 µg/ml streptomycin (ThermoFisher, #15070), 20 ng/ml EGF (Peprotech, #AF100-15), 0.5 mg/ml hydrocortisone (Sigma, #H0888), 100 ng/ml cholera toxin (Sigma, #C8052) and 10 µg/ml insulin (Sigma, #I1882)(*47*), at 37°C in a humidified incubator with 5% CO_2_. Confluent cells were passaged every 2/3 days at 1:4 dilution.

### Polyacrylamide (PAA) gel substrates

Glass-bottom 6-well dishes (#0 thickness, IBL, #220.200.020) were treated with a bind-silane solution consisting of PlusOne Bind-Silane (VWR, #17-1330-01) and acetic acid (Pancreac Quimica, #131008-1612) in absolute ethanol for 1 hr at room temperature (RT) in a fume hood. Wells were washed 3 time with ethanol and dried. PAA gels with a Young’s modulus of 12 kPa were prepared by mixing 18.8% of 40% w/v acrylamide (Bio-Rad, #1610140), 8% of 2% w/v bis-acrylamide (Bio-Rad, #1610142), 0.5% of 10% ammonium persulfate (APS, Bio-Rad, #161-0700), 0.05% of N,N,N,N’-tetramethylethylenediamine (TEMED, Sigma Aldrich, #T9281), 0.7% of FluoSpheres carboxylate modified microspheres (0.2µm, dark red, ThermoFisher, #F8807) in HEPES solution (ThermoFisher, #15630056). 22 µl of solution was placed on the treated glass well and covered with 18 mm diameter coverslip. After 1hr polymerization at RT, PBS was added to the wells and coverslips were carefully removed. Gels were washed with PBS.

### Polydimethylsiloxane (PDMS) membranes

SU8-50 master containing circular patterns of 400 µm diameter and 50 µm height was prepared using conventional photolithography. PDMS was spin-coated on the masters to a thickness lower than the height of the SU8 features (15 µm) and cured overnight at 85°C. A thick border of PDMS was left at the edges of the membranes for the handling purpose. PDMS membranes were peeled off and kept at 4°C until use.

### Collagen patterning

To pattern collagen on top of PAA gels, gels were functionalized with a solution of 1mg/ml Sulfo-SANPAH (sulfosuccunimidyl 6-(4’-azido-2’-nitrophenylamino) hexanoate, ThermoFisher, #22589) for 5 min under UV lamp (XX-15, UVP) under 365 nm wavelength. After washing off remaining Sulfo-Sanpah with sterile PBS, gels were left to air dry for 20 min inside a cell culture hood. PDMS membranes, passivated in 2% Pluronic F-127 (Sigma, #P2443) in ddH_2_O for at least 24 hours, were washed in PBS and air-dried inside a cell culture hood. PDMS membranes were placed on top of the PAA gels, and 50 µl of 0.1 mg/ml of rat tail collagen type I solution (First Link, #112296) was placed on top of the patterns. PAA gels were incubated overnight at 4°C.

### Monolayer patterning

PDMS membranes were removed from top of the polyacrylamide gels by first adding sterile PBS. Gels were washed with PBS and incubated with 200 µl of 0.1mg/ml PLL-g-PEG solution (PLL(20)-g[3.5]-PEG(2), SUSOS AG) for 30 min at 37°C. In the meantime, MCF10A cells were trypsinised and counted. Following incubation, gels were washed once with PBS and air-dried for 5 min. 50 µl of MCF10A cell suspension containing 50,000 cells, was placed on top of the gel. Cells were incubated for 1 hour for attachment, the non-attached cells were washed 3 times with PBS, and cells were incubated in DMEM-F12 media for 24 hr in 5% CO_2_ at 37°C.

### Drug treatment

After 24 hr incubation, supernatant was aspirated and fresh DMEM-F12 medium containing 4-hydroxytamoxifen (4-OHT, 100 nM, Sigma, #H7904) or equivalent amount of DMSO (1:1000, control) was added to conditionally express *HRAS*.

### Western Blot

To confirm HRAS induction with 4-OTH, total cell protein lysates were obtained by lysing cells exposed to 100 nM 4-OHT with RIPA buffer (Thermo Fisher, #89900) containing phosphatase inhibitor cocktail 1 and 2 (Sigma Aldrich, P5726 & P2850) and protease inhibitor cocktail (Sigma Aldrich, #11836170001). Protein content was quantified with BCA Protein Assay kit according to manufacturer’s instructions (Thermo Fisher, #23227) and 20 mg of protein was mixed with 2x Laemmli buffer (Sigma Aldrich, #S3401) and boiled for 5 min at 95°C. Protein were separated using MOPS buffer (Thermo Fisher, #NP0001) in NuPAGE 4-12% Bis Tris Protein Gels (Thermo Fisher, #NP0323BOX) at 150V for 70 min at RT. Proteins were transferred to a nitrocellulose membrane at 100V for 60 min at 4°C. Membranes were blocked for 30 min in 5% milk in TBST, followed by overnight incubation with primary antibodies diluted in 2.5% milk in TBST. Primary antibodies included rabbit phospho p44/42 MAPK (ERK1/2) (1:2000, Cell Signalling, #4370S), rabbit p44/42 MAPK (Erk1/2) (1:2000, Cell Signalling, #4695S), mouse E-cadherin (1:1000, BD, #610181). Primary antibodies were washed 3 times with TBST, followed by 1 hour incubation with secondary antibodies diluted in 2.5% milk in TBST (1:5000, goat anti-rabbit HRP, or goat anti-mouse HRP, DAKO, #P0448 & #P0447). Membranes were washed 3 times with TBST and exposed to HRP substrate (Immobilon Crescendo, Millipore, #WBLUR0100) for chemiluminescence detection using ChemiDoc™ MP Imaging System (Bio-Rad).

### Time-lapse microscopy

Multidimensional acquisitions were performed on an automated inverted microscope (Nikon Ti2 Eclipse, Nikon) using 20x objective (Nikon CFI Plan Apo 20X/0.75 Ph DM). Microscope was equipped with thermal, CO_2_ and humidity control, and controlled using NIS software and perfect focus system. Images were obtained every 15 min over 50 hours. Up to 15 independent patterns were imaged in parallel using a motorized XY stage.

### Traction force microscopy

Traction forces were computed from hydrogel displacements through a custom-made software developed in the laboratory of Xavier Trepat, which is based on a Fourier-transform algorithm for elastic hydrogel substrates having finite thickness(*48*). Gel displacements between any experimental time point and a reference image obtained after cell trypsinization were computed by using a custom-made particle imaging velocimetry code developed in the laboratory of Xavier Trepat by using 32-pixel resolution and overlap of 0.5.

### Monolayer stress microscopy

Inter-cellular and intra-cellular stresses in non-transformed MCF10A monolayers were computed via Monolayer Stress Microscopy(*31*) via a custom software implemented via the custom FEM-platform EMBRYO developed in the laboratory of José Muñoz. Briefly, the forces exerted by the elastic hydrogel substrate on the MCF10A epithelium (as a reaction to the traction force field transferred by the epithelium to the substrate) are equilibrated by the tensorial stress state within the epithelium. A necessary condition for the application of this technique is that epithelia maintain their monolayer architecture, a hypothesis only valid for non-transformed MCF10A monolayers in this study.

### Immunofluorescence

Cells were fixed with 4% paraformaldehyde (PFA, Santa Cruz, #sc-281692) for 10 min and washed with PBS. Samples were incubated with block buffer containing 1% bovine serum albumin (BSA, Sigma, #A7906) and 0.3% Triton-X100 (Sigma, #T8787) in PBS at RT for 1 hr. Primary antibodies (mouse E-cadherin, 1:1500, BD Biosciences, #610181; mouse integrin ß1, 1:250, Abcam, ab30394; rabbit pMLC2, 1:50, Cell Signalling, #3671S) were diluted in block buffer and incubated on top of samples overnight at 4°C. Subsequently, samples were incubated with secondary antibodies (FITC anti-mouse, 1:1000, Jackson Immuno Research, #715-545-150; AlexaFluor564 anti-rabbit, 1:500, Thermo Fisher, #A11035) for 2hr at RT. F-actin was stained by incubating for 30 min with phalloidin-iFluor594 cytoPainter (1:2000, Abcam, #ab176757) or Phalloidin-Atto 647N (1:1000, Sigma-Aldrich, #65906) at RT. In between steps, samples were washed with wash buffer (0.05% Tween-20 (Sigma, #P9416) in PBS). Samples were covered with Fluoroshield mounting medium containing DAPI (Sigma, #F6057) and stored at 4°C until imaging.

### Microscopy

Fluorescent images of the patterns were acquired with an inverted microscope (Nikon Ti2 Eclipse, Nikon) with an objective 20x/0.75 (Nikon CFI Plan Apo 20X/0.75 Ph DM). Confocal images were taken using inverted confocal microscope Axio Observer 7 (Spectral Detection Zeiss LSM 800) using 40x/1.3 Oil DIC M27 or 63x/1.4 Oil DIC M27 objectives, with ZEN 2.3 imaging software. For integrin imaging, Zeiss Axiovert 200M microscope was used with 20x and 40x objective. For pMLC2 imaging, Zeiss LSM 780 microscope was used with 20x/0.8 M27 Plan-Apochromat and 40x/1.20 W Korr M27 C-Apochromat objectives using ZEN 2.1 SP3 software.

### Nanoindentation

Monolayer stiffness (Young’s modulus) was measured by means of the Piuma Nanoindenter (from Optics 11) fitted with a cantilever having stiffness of 0.05 N/m and spherical tip with a radius of 10 µm. Four measurements were taken from 3 different samples. The Young’s modulus of tissues before oncogene induction resulted to be 1.363±0.504 kPa (Mean±SD).

### Image analysis

In order to detect the physical properties of the epithelial monolayer from fluorescent images a pipeline was created in CellProfiler(*49*) and followed by post processing of the images and data in custom-made automatic workflow in MATLAB (License Number 284992). Images of nuclei and F-actin were used to detect both the individual nuclei and cell borders within monolayers. The intensity of the images was rescaled to the full range of the intensity histograms (minimum 0, maximum 1) and uneven illumination was corrected by subtracting a spline illumination function. Nuclei were segmented using the adaptive Otsu three-class thresholding with the middle intensity class assigned as the background. To improve detection, we optimized minimum diameter (16-pixel unit) and threshold correction factor (0.8). Clumped objects were distinguished by intensity and cell outlines by propagation method. The precision of nuclei detection was assessed by comparing the outcomes of the pipeline with manual nuclei detection in ImageJ. Features extracted from the image processing included cell ID, nuclei centres coordinates, areas and perimeters. Using custom-made workflow in MATLAB (License Number 284992), we calculated cell and nuclei shape indices and removed outliers (based on surface areas, shape index, nucleus areas).

### E-cadherin analysis

To measure E-cadherin intensity a line was drawn using Fiji ImageJ(*50*) (version 1.53c) between two nuclei. The intensity range was then normalized by subtracting the 1^st^ percentile and dividing by 99^th^ percentile. The junctional intensity was extracted, and the ratio between junctional and cytoplasmic intensity was calculated by dividing the junctional value by average of the cytoplasmic value. **Integrin analysis**. To measure integrin intensity a global intensity value of the total image was obtained with ImageJ. **pMLC2 analysis**. Fluorescence intensity was determined using Fiji. Rectangular shape (531.37×290 µm) was placed from one end of the image to other encompassing the middle part of the pattern and the average intensity was measured. The background intensity was removed by calculating intensity in area outside of the pattern for each image. Data from three independent patterns were presented as mean±standard error.

### Data analysis

To perform statistical analysis GraphPad Prism (version 9.0.0) was used. Data distribution was assessed using D’Agostino and Pearson omnibus normality test. Data from all conditions had to pass the normality test to be included in parametric testing. For non-parametric data: a) two groups – Mann Whitney test; b) more than two groups – Kruskal-Wallis statistic test used with Dunn’s Multiple Comparison Test; c) two groups over long time-course – 2-way ANOVA with Bonferroni post-test; d) one condition over long time-course – Friedman test with Dunn’s multiple comparison test. P-value below 0.5 indicated statistical significance (*p<0.05, **p<0.01, ***p<0.001).

### Computational model

The evolution of the net radial-traction components is modelled by resorting to a two-dimensional Finite Element (FE) model of the flat tissue. The flat tissue is represented by a circular domain Ω with radius *R*, with two distinct subdomains: subdomain Ω_1_ subjected to a constant baseline contractile force 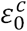 (pre-strain) and subdomain Ω_2_ subjected to an additional active contractile strain *ε*^*c*^, for a total contractile strain of 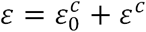 (Fig.4B). The elastic domains Ω_1_ and Ω_2_ develop local tension as a consequence of these prescribed strains. Moreover, each domain is subjected to a specific degree of elastic adhesion with the underlying substrate, which is modelled as a set of nodal locations fixed in time. Weakening of cell-matrix adhesion is simulated by applying a reduction factor α to the cell-matrix adhesion constant κ. The subdomains Ω_1_ and Ω_2_ are assumed to have linear elastic behavior and tissue motion is determined in the approximation of quasi-static equilibrium – there, the active contractility 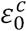 and the baseline pre-strain 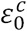 equilibrate the passive elastic forces within the tissue and the adhesion forces with the substrate. We then used experimental data to set the model’s parameters and Cauchy’s equation for elastic continua to determine model’s deformations in each of the subdomains Ω_1_ and Ω_2_ (Supplementary Text 2).

## Acknowledgements

This work was supported by the Spanish Ministry MICINN /FEDER (grants BFU2016-75101-P and RYC-2014-15559 to V.C., DPI2016-74929-R to J.J.M., BES-2017-081337 to G.F., BES-2013-062633 to M.U.); the Generalitat de Catalunya (grant 2017SGR1278 to J.J.M.); the IBEC-ICMS Exchange Program fund to A.N. for nanoindentation measurements; H.K.M was supported by a CRUK/EPSRC Multi-disciplinary Project Award (C1529/A23335) and the MRC/UCL Laboratory for Molecular Cell Biology (MC_CF12266). The Spanish Ministry (MICINN) and its funding programs (Severo Ochoa, FPI, Excelencia) along with the CERCA Program of the Generalitat de Catalunya support research at the IBEC. The authors would also like to thank: Carlos Pérez-González (IBEC); Víctor González-Tarragó (IBEC); Manuel Gómez González (IBEC); Prof Bouten (TU/e, ICMS); and, Janine Grolleman (TU/e, ICMS) for helpful technical advice and support.

## Author contributions

A.N. and V.C. designed experiments; A.N. performed all experiments with contributions from S.D., G.F. and H.M.; A.N., M.U. and H.M. designed protocols; J.M. designed and implemented the computational model; B.B. and H.M. provided the cellular model; X.T. provided custom software for Traction Force Microscopy; A.N., J.M., S.D. and V.C. performed data analysis; A.N., J.M., S.D., X.T., H.M., B.B. and V.C. interpreted data; V.C. conceived the study, secured the funding and supervised the project. A.N., J.M. and V.C. wrote the manuscript, which all authors reviewed and edited.

## Data availability

All data supporting the findings of this study are available from the corresponding author upon reasonable request.

## Code availability

Custom code may be made available from the corresponding author upon reasonable request.

