## Supplementary Materials for "Oncogenic *RAS* instructs morphological transformation of human epithelia via differential tissue mechanics"

**This PDF file includes:**

Supplementary Text 1 to 2
Figures S1 to S10
Table S1
Captions to Movies S1 to S4

**Other Supplementary Materials for this manuscript include the following:**

Movies S1 to S4

**Supplementary Text 1**

Topology of the traction-force field.

To analyse the distribution and orientation of the traction forces among different epithelial subdomains, we subdivided the epithelial subdomain in four elliptical quadrants $Q_1, \ldots, Q_4$. Elliptical quadrants of each epithelial domain at each time-point were determined as follows. Firstly, the elliptical hull is identified that best approximates the epithelial domain at each time-point – the elliptical hull is defined as the elliptical envelope/hull of the epithelial domain having the same normalized second central moments as the epithelial domain which it approximates. Secondly, the reference system *C-ab* is identified that is centred in the centroid $\boldsymbol{C}$ (black point in Fig. S5,S7 A-B) of the epithelial domain and that has its horizontal axis *a* (horizontal white arrow in Fig. S5,S7 A-B) oriented along the major axis of the elliptical hull and its vertical axis *b* oriented along the minor axis (vertical white arrow Fig. S5,S7 A-B) of the elliptical hull. The elliptical quadrants are then determined by rotating the *C-ab* reference system by 45 degrees contra clockwise (Fig. S5,S7 A-B), so that $Q_1$ and $Q_3$ subdomains are astride the *a*-axis (major axis of the elliptical hull) whereas $Q_2$ and $Q_4$ subdomains are astride the minor $b$-axis (minor axis of the elliptical hull). Next, we computed resultant traction-force vector $\vec{\boldsymbol{T}}_{\boldsymbol{Q}}$ obtained by vectorially summing all traction-force vectors $\vec{\boldsymbol{T}}$ in each elliptical quadrant $Q_i$ for each epithelium at each time point. then magnitudes and angles are computed. Next, we adopted a reference system **C**-*ab* centred in the centroid **C** of the epithelial domain and oriented along the major and minor axes (respectively *a* and *b*) of the elliptical hull at each time point (Fig. S5,S7 A-B). We then calculated for each time-point of the time-evolution of each epithelial domain: i) the modulus $\|\vec{\boldsymbol{T}}_{\boldsymbol{Q}}\|$ of each of the for resultants (i.e. the Euclidean norm or length of the vector); and, ii) the angle (in the interval $[-\pi, +\pi]$) that each resultant vector $\vec{\boldsymbol{T}}_{\boldsymbol{Q}_i}$ forms with its respective unit directions (Fig. S5,S7 A-B). Angles are obtained from the scalar product between $\vec{\boldsymbol{T}}_{\boldsymbol{Q}_i}$and its respective $\boldsymbol{e}_i$. It is worth observing that $\boldsymbol{e}_1$ and $\boldsymbol{e}_2$ are the unit directions antiparallel to the *a*-axis and *b*-axis of the **C**-*ab* reference system at each time point, the uni-directions $\boldsymbol{e}_3$ and $\boldsymbol{e}_4$ being parallel axes *a* and *b* respectively. Accordingly, a zero-degree angle between $\vec{\boldsymbol{T}}_{\boldsymbol{Q}_i}$ and $\boldsymbol{e}_i$ meant that the resultant vector $\vec{\boldsymbol{T}}_{\boldsymbol{Q}_i}$ was parallel to $\boldsymbol{e}_i$ and, thus, pointing towards the origin $\boldsymbol{C}$ of the reference system *C-ab* (i.e. towards the epithelial domain's centroid). Finally, we computed calculated for each time-point of the time-evolution of each epithelial domain the traction-force resultant from opposite elliptical quadrants – i.e. the force dipoles $\vec{\boldsymbol{T}}_{13} = \vec{\boldsymbol{T}}_{\boldsymbol{Q}_1} + \vec{\boldsymbol{T}}_{\boldsymbol{Q}_3}$ and $\vec{\boldsymbol{T}}_{24} = \vec{\boldsymbol{T}}_{\boldsymbol{Q}_2} + \vec{\boldsymbol{T}}_{\boldsymbol{Q}_4}$ (simply referred to as $\vec{\boldsymbol{T}}_{DIPOLES}$) along with their respective magnitudes $\|\vec{\boldsymbol{T}}_{13}\|$ and $\|\vec{\boldsymbol{T}}_{24}\|$.

We observed that: i) the time-evolution of $\|\vec{\boldsymbol{T}}_{\boldsymbol{Q}_i}\|$ in each elliptical quadrant $Q_i$ (Fig. S5,S7) followed a trend similar to that of traction force magnitudes respectively for non-transformed (Fig. 2 F), *HRAS*-transformed (Fig. 2 G) and *KRAS*-transformed epithelia (Fig. S6 N); ii) vectors $\vec{\boldsymbol{T}}_{\boldsymbol{Q}_i}$ in the periphery of epithelial domains are centripetal (i.e. pointed in average towards the domain's centre) for both normal and *RAS*-transformed tissues (Fig S5,S7 M) – this is evident from both non-transformed and *RAS*-transformed angle trends evolving in time within a narrow span of approximately ±20 degrees of the epithelial domain's centroid; iii) vectors $\vec{\boldsymbol{T}}_{\boldsymbol{Q}_i}$ in the central part of the epithelial domains lose their centripetal character and become more randomly oriented (Fig S5,S7 H) – this is evident from both non-transformed and *RAS*-transformed angle trends evolving in time within a much wider span of approximately ±45 degrees of the epithelial domain's centroid; and, iv) epithelial domains start losing their constant circularity (Fig. 1 H and Fig. S6 I) and aspect ratio (Fig. 1 I and Fig. S6 J) concomitantly to tractions dipole across the major diameter of the elliptical hull (Fig. 1 G and Fig. S6 H) starting to increase in magnitude more than the dipole across the minor diameter of the epithelial domain (Fig. S5,S7 N-O).

**Supplementary Text 2**

In silico model of RAS-driven mechanical instability.

The evolution of the net radial-traction components is modelled by resorting to a two-dimensional Finite Element (FE) model of the flat tissue. The flat tissue is represented by a circular domain Ω with radius *R*, with two distinct subdomains: subdomain $\Omega_1$ subjected to a constant baseline contractile force $\varepsilon_0^c$ (pre-strain) and subdomain $\Omega_2$ subjected to an additional active contractile strain $\varepsilon^c$, for a total contractile strain of $\varepsilon = \varepsilon_0^c + \varepsilon^c$ (Fig. 4 B). The elastic domains $\Omega_1$ and $\Omega_2$ develop local tension because of these prescribed strains. Moreover, each domain is subjected to a specific degree of elastic adhesion with the underlying substrate, which is modelled as a set of nodal locations fixed in time. Weakening of cell-matrix adhesion is simulated by applying a reduction factor α to the cell-matrix adhesion constant κ. The subdomains $\Omega_1$ and $\Omega_2$ are assumed to have linear elastic behaviour and tissue motion is determined in the approximation of quasi-static equilibrium – there, the active contractility $\varepsilon_0^c$ and the baseline pre-strain $\varepsilon_0^c$ equilibrate the passive elastic forces within the tissue and the adhesion forces with the substrate.

Model's deformations in each subdomain $\Omega_1$ and $\Omega_2$ were determined via Cauchy's equation

$$\nabla \cdot \boldsymbol{\sigma} + \boldsymbol{f} = \mathbf{0}, \qquad (1)$$

where:

$\boldsymbol{\sigma}$ is the elastic stress tensor given by $\boldsymbol{\sigma} = \lambda\, trace(\boldsymbol{\varepsilon}^e) + 2\mu\boldsymbol{\varepsilon}^e$;
$\boldsymbol{\varepsilon}^e$ is the elastic strain, with $\boldsymbol{\varepsilon}^e = \boldsymbol{\varepsilon} - \varepsilon_0^c \boldsymbol{I}$ in $\Omega_1$ and $\boldsymbol{\varepsilon}^e = \boldsymbol{\varepsilon} - (\varepsilon_0^c + \varepsilon^c)\boldsymbol{I}$ in $\Omega_2$;
$\lambda = E\nu/((1+\nu)(1-2\nu))$. and $\mu = E/(2(1+\nu))$ are the Lamé elasticity constants;
$E$ is the Young modulus and ν the Poisson ratio of both subdomains $\Omega_1$ and $\Omega_2$;
$\boldsymbol{f} = \alpha k \boldsymbol{u}$ is the adhesion force between the tissue and the substrate;
$\boldsymbol{u}$ is the tissue displacement and $u_i$ its components;
$\varepsilon_{ij} = 0.5(\partial_i u_j + \partial_j u_i)$ is the total (observable) strain; and,
$\alpha$ is the weakening factor of the tissue adhesion forces with the substrate ($\alpha = 1$ in $\Omega_1$).

We use the Finite Element Method to turn equation (1) into a linear system of equations

$$(\mathbf{K} + \mathbf{K}_s)\mathbf{u} = \mathbf{f} \qquad (2)$$

where $\mathbf{K}$ and $\mathbf{K}_s$ are the stiffness matrices corresponding to the elastic and adhesive contributions respectively. Matrices $\mathbf{K}$ and $\mathbf{K}_s$ are computed via the formulas

$\mathbf{K} = \int_\Omega \mathbf{B}^T\mathbf{D}\mathbf{B}\, d\Omega$ and $\mathbf{K}_s = \int_\Omega \alpha k \mathbf{N}^T\mathbf{N} d\Omega$

where:

$\mathbf{B}$ is the deformation matrix such that $\boldsymbol{\varepsilon} = \mathbf{Bu}$;
$\mathbf{D}$ is the elasticity matrix that depends on $E$ and ν; and,
$\mathbf{N}$ is a matrix containing the Finite Element interpolation functions.

The force vector **f** in equation (2) represents the resulting contractile force due to active contractile strain $\varepsilon^c$ and/or baseline pre-strain $\varepsilon_0^c$. The time evolution of $\alpha, \varepsilon^c$and $\varepsilon_0^c$.are given in Supplemental Table 1. Vector **f** is determined from the assembling of vectoral contributions $\mathbf{f_1}$ and $\mathbf{f_2}$ respectively from the subdomains $\Omega_1$ and $\Omega_2$, with:

$\mathbf{f_1} = \varepsilon_0^c \int_{\Omega_1} \mathbf{B}^{\boldsymbol{T}} \mathbf{D} \{1\ 1\ 0\}^{\boldsymbol{T}}\, d\Omega$ and $\mathbf{f_2} = (\varepsilon_0^c + \varepsilon^c) \int_{\Omega_2} \mathbf{B}^{\boldsymbol{T}} \mathbf{D} \{1\ 1\ 0\}^{\boldsymbol{T}}\, d\Omega$.

We then used experimental data to set: i) the bulk stiffness of the epithelium as having a value of 1kPa prior to *HRAS* activation – this is based on nanoindentation measurements of MCF10A epithelia having an *in vitro* stiffness of 1.36kPa ±0.5kPa (see Materials & Methods). Furthermore, for the sake of simplicity, we also set: ii) the Poisson-Ration of the tissue elasticity to ν=0.46 (quasi-incompressibility); iii) the drop in adhesion with the substrate (Fig. 6 B,E) to a value of 70% from baseline value – this corresponds to the average drop in the expression of the collagen receptor integrin β1 within 24 hours from HRAS activation (Fig. 4 E-F); and, iv) a linear increase in local contractile strain $\varepsilon = \varepsilon_0^c + \varepsilon^c$ (Fig. 6 B,F) to a value approximately double that of the homeostatic baseline (Table S1) – this is based on combined pMLC2 fluorescence intensity from bottom and top layers of the *RAS*-transformed bilayer (Fig. 5 F,H) that approximately amounts to double that of non-transformed monolayers (Fig. 5 C).

It is worth noticing that a local reduction in the area of the confined epithelium induced by pre-strains and active contractility of each finite element contributes in a linear manner to the global reduction in area of the whole epithelium. However, the intensity of traction forces transferred by each finite element to the substrate strongly depends on the topological distribution of active tension within the tissue, which differs between the three topological mechanisms tested (Fig. 6 G,J,M). That is because contractility along shared edges of neighbouring finite element of the discretized epithelium (Fig. 6 C-D) partly tends to cancel each other out and partly gets transferred to the substrate (Fig. 6 B).

**Fig. S1.**

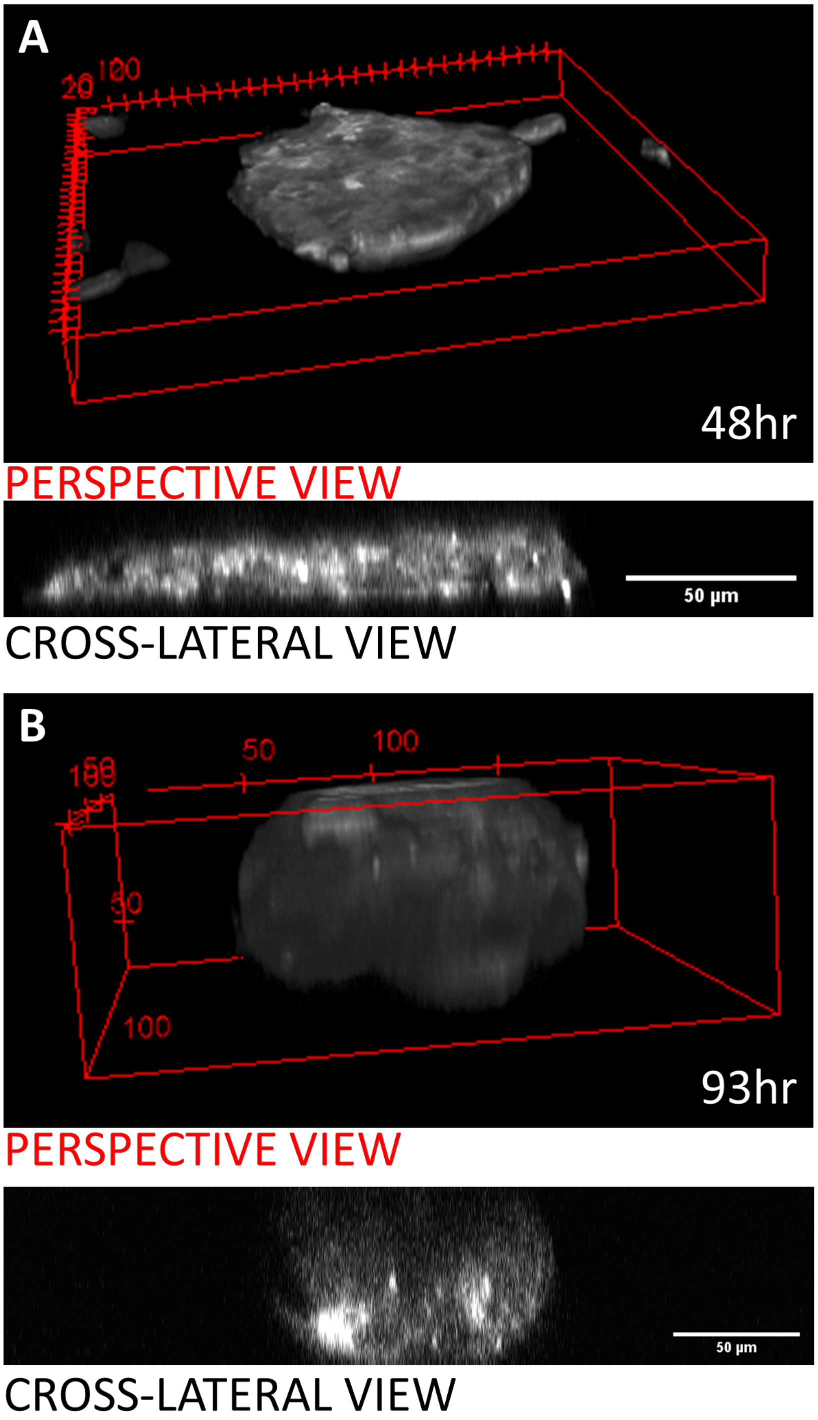


**The HRAS cell mass.**
**A-B)** Confocal microscopy reconstruction of HRAS-transformed 3D cell mass at t=48 (A) and t=93 hours (B) both in perspective and cross-lateral view.

**Fig. S2.**

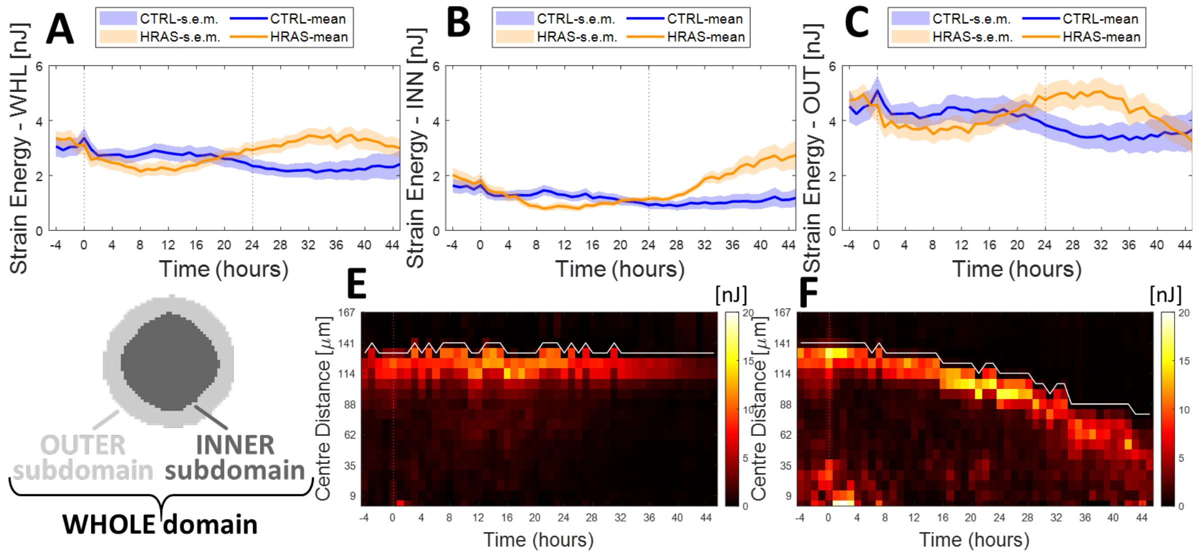


**Strain Energy. A-C)** Time-evolution (Mean±S.E.M.) of the elastic strain energy intensity actively transferred by the epithelium to the elastic gel substrate (Mean±S.E.M). Elastic strain energy was defined as one-half the scalar product between cellular traction and the gel displacement that it generates. Statistics over 15 non-transformed epithelia and 16 HRAS-transformed epithelia from at least 4 independent experiment repeats. Strain energy trends in: **A)** the whole island's domain; **B)** inner island's subdomain; and, **C)** the outer island's subdomain. **D)** Schematic representing the whole epithelial domain along with its outer and inner subdomains. **E-F)** Kymographs of the strain energies: **E)** for a representative control MCF10A circular epithelium; and, **F)** for a representative *HRAS*-transformed MCF10A circular epithelium. White lines represent the average evolution of the edge of the island in time, the centre of the island co-localizing with the bottom of the graph.

**Fig. S3.**

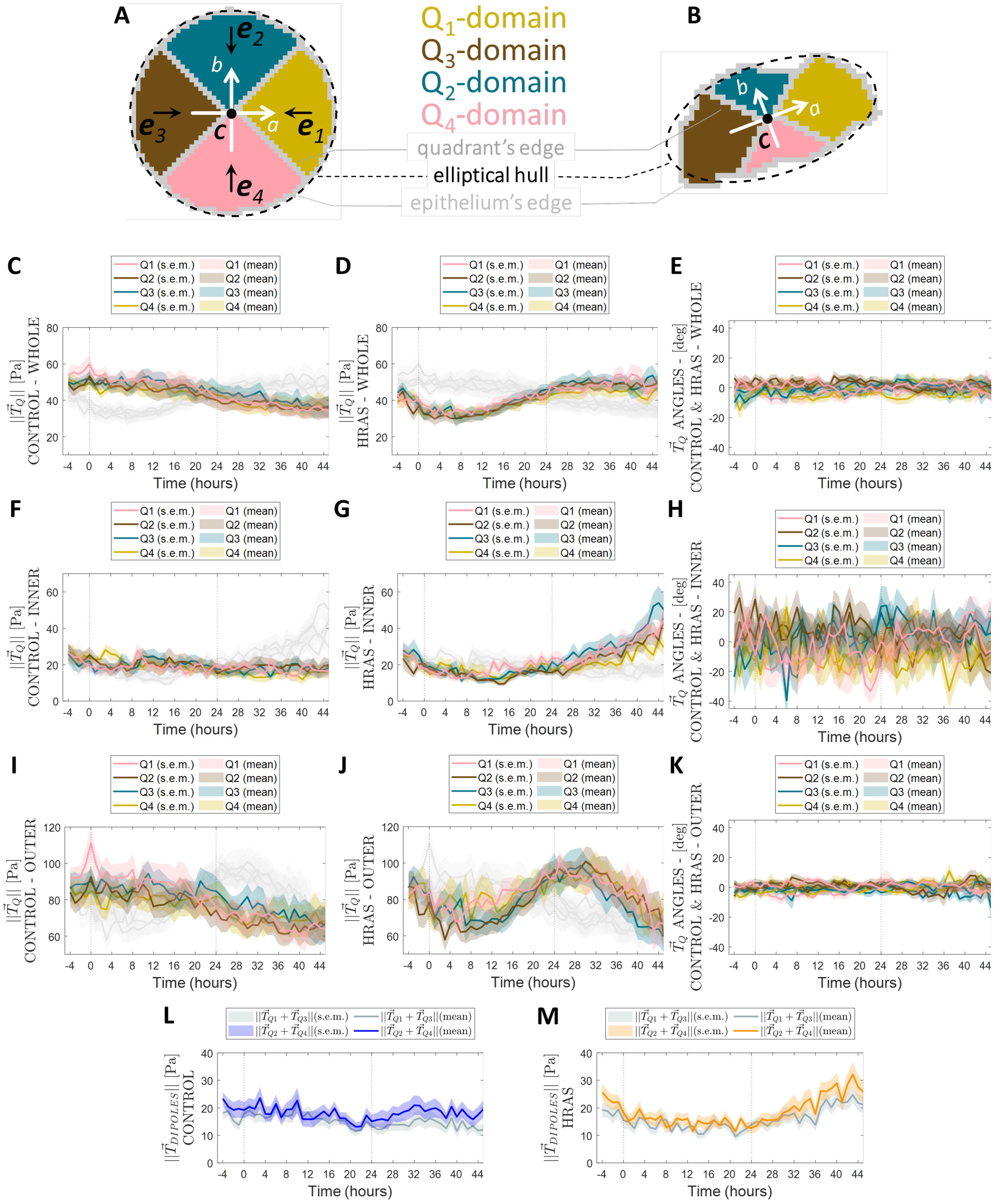


**Cell traction forces in HRAS-transformed MCF10A monolayer.**

**A-B)** Cartoons showing the subdivision of the whole epithelial domain (grey circle) into four **elliptical quadrants** $Q_1, \ldots, Q_4$ respectively: **A)** at a representative early time point; and, **B)** at a representative later time point. The intersection between each elliptical quadrant $Q_i$ with the inner and outer subdomain of the whole epithelial domain are referred to as the **inner and outer elliptical quadrants**. **C-M)** Time-evolution (Mean±S.E.M.) of the magnitude and angle of the resultant traction-force vector $\vec{\boldsymbol{T}}_{\boldsymbol{Q}}$ obtained by vectorially summing all traction-force vectors $\vec{\boldsymbol{T}}$ from each elliptical quadrant $Q_i$ – traction force vectors are added vectorially in each quadrant for each epithelium at each time point, then magnitudes and angles are computed. Magnitudes are intended in the sense of $\|\vec{\boldsymbol{T}}_{\boldsymbol{Q}}\|$ (the Euclidean norm or length of the vector) whereas the angle is intended as the trigonometric angle (in the interval $[-\pi, +\pi]$) formed by each vector $\vec{\boldsymbol{T}}_{\boldsymbol{Q}_i}$ with the unit directions $\boldsymbol{e}_i$. Time-evolution (Mean±S.E.M.) of $\|\vec{\boldsymbol{T}}_{\boldsymbol{Q}}\|$ for non-transformed epithelia in the whole (**C**), inner (**F**) and outer (**I**) elliptical quadrants and *HRAS*-transformed epithelia in the whole (**D**), inner (**G**) and outer (**J**) elliptical quadrants. **E**, **H**, **K)** Time-evolution (Mean±S.E.M.) of the angle between vector $\vec{\boldsymbol{T}}_{\boldsymbol{Q}_i}$ with the unit directions $\boldsymbol{e}_i$ in each elliptical quadrant $Q_i$ for non-transformed and *HRAS*-transformed epithelia in the whole (**E**), inner (**H**) and outer (**K**) elliptical quadrants. Time-evolution (Mean±S.E.M.) of force-resultant magnitudes from opposite elliptical quadrants (see subpanels A-B) – i.e. $\|\vec{\boldsymbol{T}}_{\boldsymbol{Q}_1} + \vec{\boldsymbol{T}}_{\boldsymbol{Q}_3}\|$ **and** $\|\vec{\boldsymbol{T}}_{\boldsymbol{Q}_2} + \vec{\boldsymbol{T}}_{\boldsymbol{Q}_4}\|$ (also simply referred to as $\vec{\boldsymbol{T}}_{DIPOLES}$): **L)** for non-transformed circular epithelia; and, **M)** for *HRAS*-transformed circular epithelia.

**Fig. S4.**

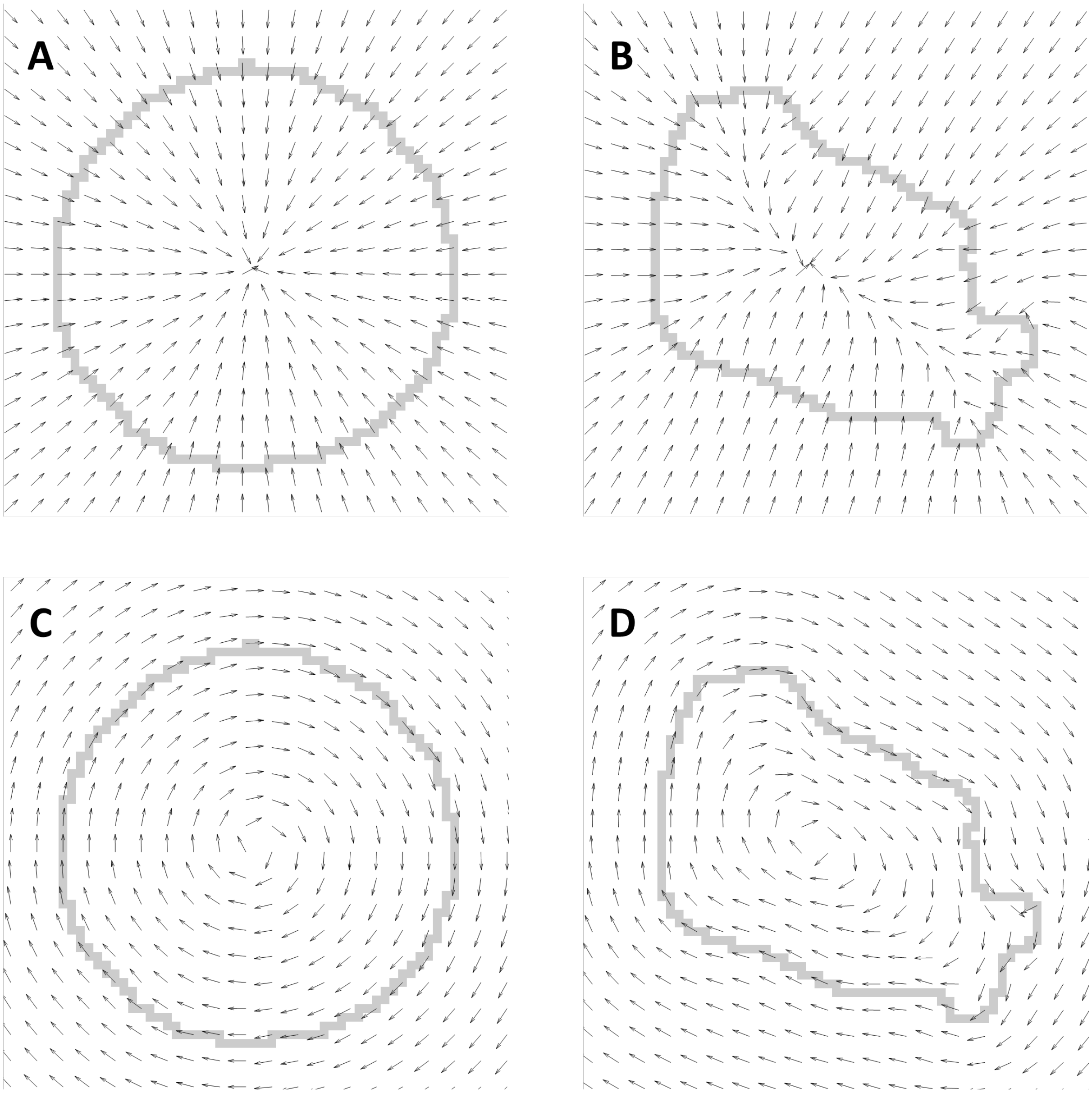


**Vector fields of normal and tangent directions. A-D)** Representative diagrams showing the mutually orthogonal fields of unit directions (black arrows) that are: **A-B)** perpendicular to the epithelial domain's edge (grey) in an earlier (A) and later (B) stage of epithelial evolution; and, **C-D)** tangential to the epithelial domain's edge (grey) in an earlier (C) and later (D) stage of epithelial evolution.

Fig. S5.

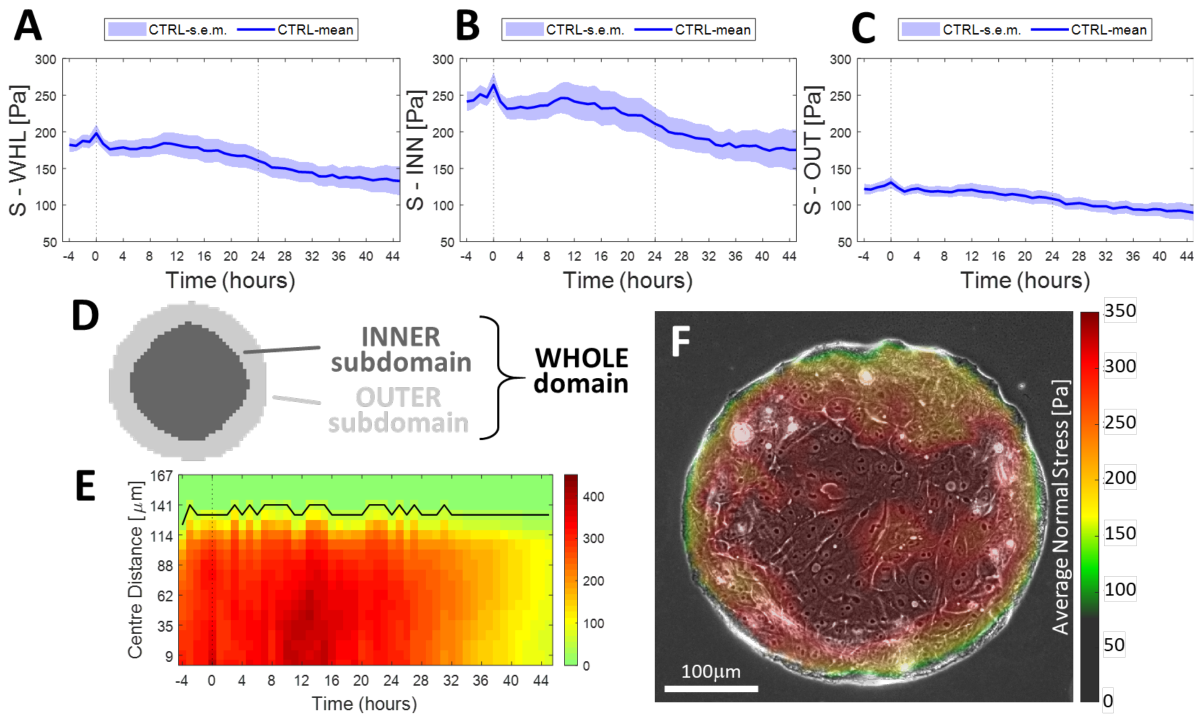


**Average normal stress within the non-transformed MCF10A monolayer.**

Time-evolution (Mean±S.E.M.) of average normal stress (Mean±S.E.M.): **A)** in the whole island's domain; **B)** in the inner island's subdomain; and, **C)** in the outer island's subdomain. Statistics over 15 non-transformed epithelia from at least 4 independent experiment repeats. **D)** Schematic representing the whole epithelial domain along with its outer and inner subdomains. **E)** Kymograph of the average normal stress for a representative control MCF10A circular epithelium. Black line represents the average evolution of the edge of the island in time, the centre of the island co-localizing with the bottom of the graph. **F)** Average normal stress within the epithelial monolayer (red is tension and grey is compression, according to the colour bar).

**Fig. S6.**

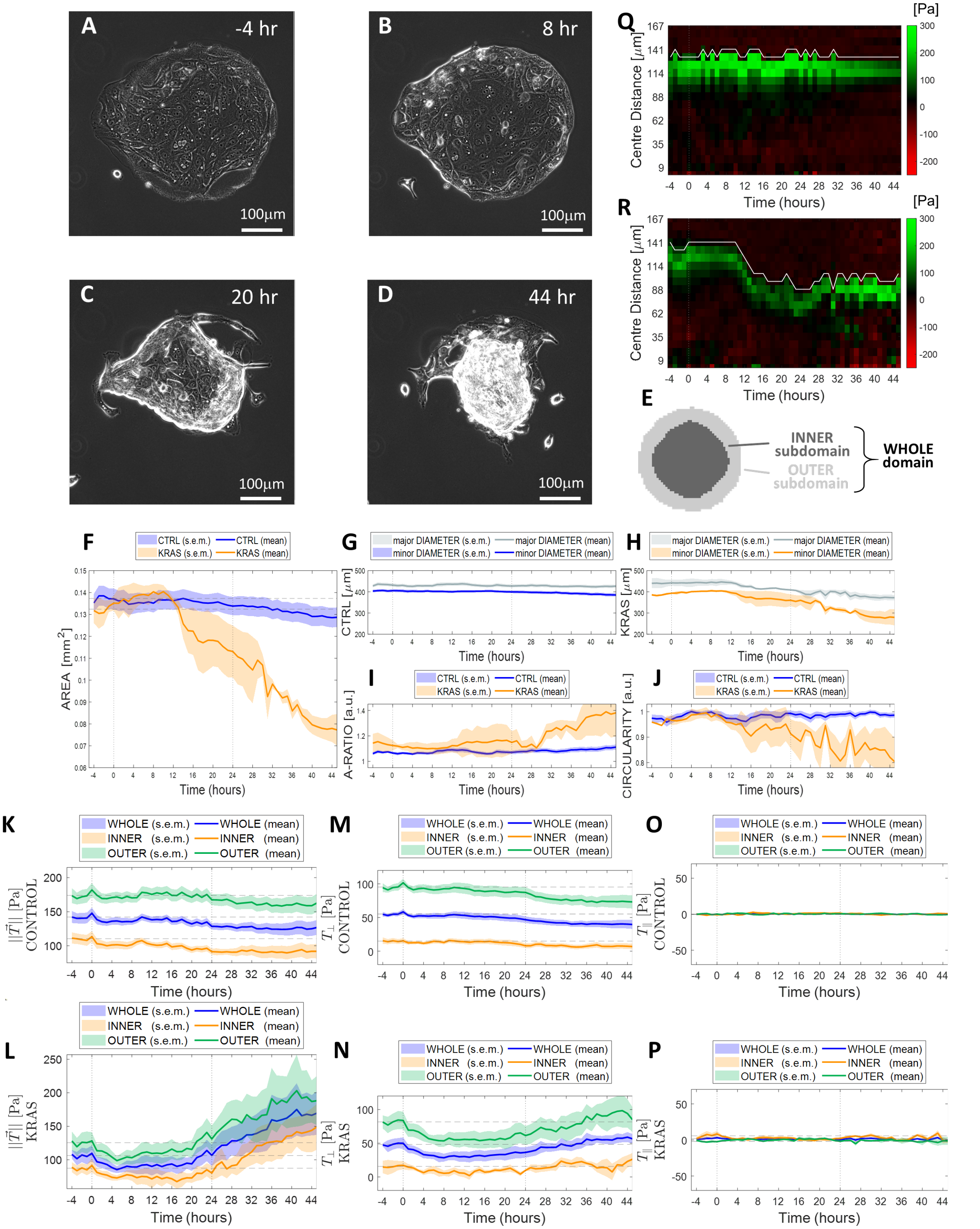

**Morpho-mechanical evolution of KRAS-transformed MCF10A monolayers. A-D)** Phase contrast time-lapse of a *KRAS*-transformed MCF10A monolayer (imaging starts at t=-4 hours and *KRAS*-activation is induced at t=0 hours; scale bar 200 μm). **E)** Schematic representing the whole epithelial domain along with its outer and inner subdomains. Time-evolution of the surface area (**F**), the major and minor diameters of the epithelia domain in the case of non-transformed (**G**) and *KRAS* -transformed (**H**) tissues; epithelial domain's aspect ratio (**I**) and the epithelial domain's circularity (**J**). Time evolution (Mean±S.E.M.) of magnitudes (**L**) and components of the traction-force field (**N,P**) in the whole epithelial domain (blue) as well as in the inner (orange) and outer (green) epithelial subdomains. **K-L)** Time evolution of the average Traction-field ±magnitude is computed as $\|\vec{T}\|$, the Euclidean norm (length) of the traction-force vectors of the field, for both: **K)** non-transformed epithelia; and, **L)** *KRAS* -transformed epithelia. **M-P)** Traction-field components are computed with their own sign by projecting traction-force vectors (at each time point and in each location of the epithelial domain) along the perpendicular and tangential directions respectively (Suppl. Fig. 3). Time evolution of the average traction force components: **M-N)** perpendicular to the island's edge ($T_\perp$) for **M)** non-transformed and **N)** *KRAS*-transformed epithelia; **O-P)** tangential to the island's edge ($T_\parallel$) for **O)** non-transformed and **P)** *KRAS* -transformed epithelia. **Q-R)** Kymographs of the perpendicular component of the traction field $T_\perp$: **Q)** for a representative non-transformed MCF10A epithelium; and, **R)** for a representative *KRAS* -transformed MCF10A epithelium. White lines represent the average evolution of the edge of the island in time, the centre of the island co-localizing with the bottom of the graph. A negative component $T_\perp$ (red) means that the corresponding traction-force vector is in oriented towards the exterior of the epithelial domain's edge, whereas a positive component $T_\perp$ (green) is indicative of traction-force orientation toward the interior of the epithelial domain's edge.

**Fig. S7.**

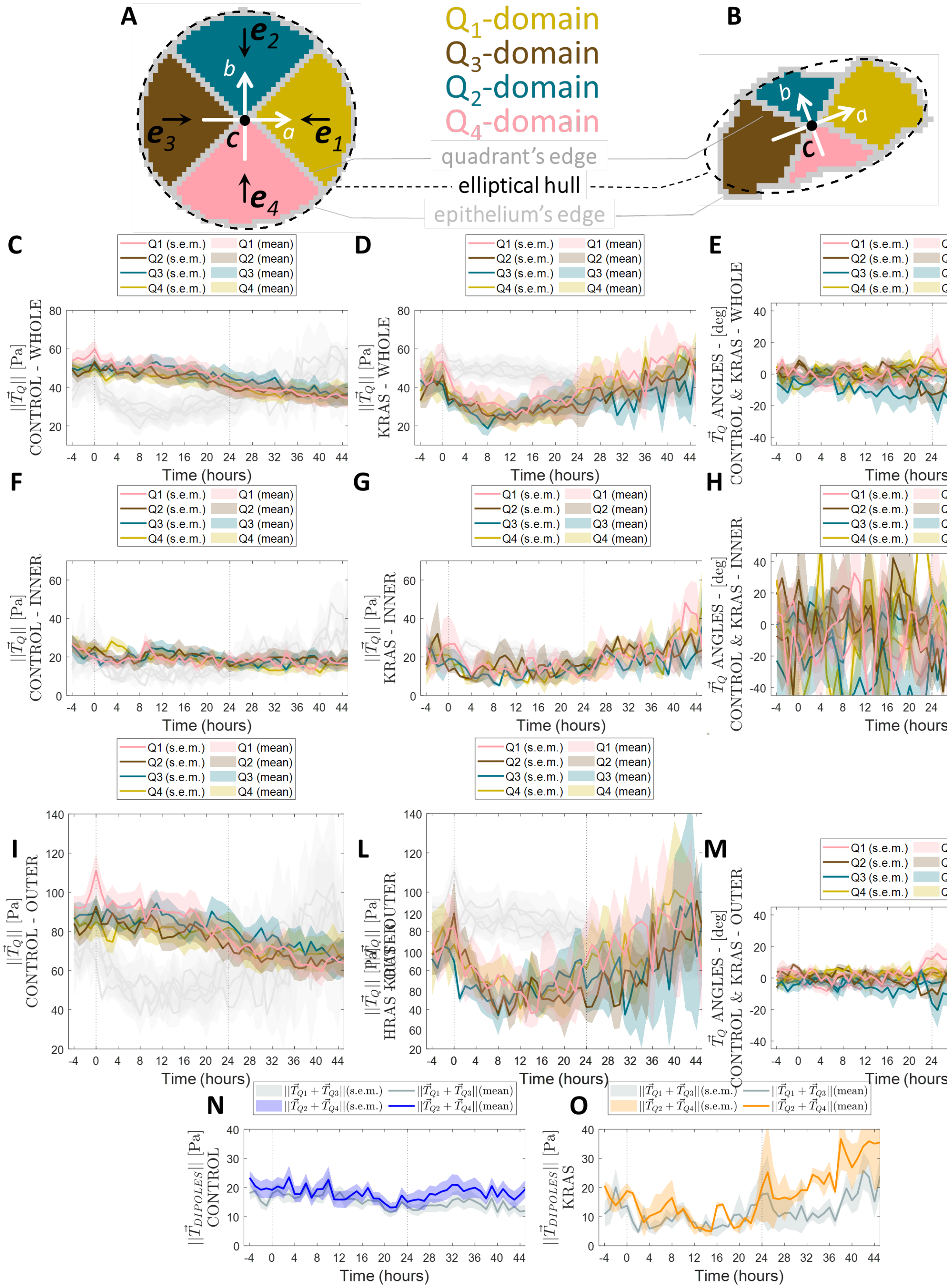


**Cell traction forces in *KRAS*-transformed MCF10A monolayer.**

**A-B)** Cartoons showing the subdivision of the whole epithelial domain (grey circle) into four **elliptical quadrants** $Q_1, \ldots, Q_4$ respectively: **A)** at a representative early time point; and, **B)** at a representative later time point. The intersection between each elliptical quadrant $Q_i$ with the inner and outer subdomain of the whole epithelial domain are respectively henceforth referred to as the **inner and**

**outer elliptical quadrants**. **C-M)**. Time-evolution (Mean±S.E.M.)of the magnitude and angle of the resultant traction-force vector $\vec{\boldsymbol{T}}_{\boldsymbol{Q}}$ obtained by vectorially summing all traction-force vectors $\vec{\boldsymbol{T}}$ from each elliptical quadrant $Q_i$ – traction force vectors are added vectorially in each quadrant for each epithelium at each time point, then magnitudes and angles are computed. Magnitudes (**C,D,F,G,I,L**) are intended in the sense of $\|\vec{\boldsymbol{T}}_{\boldsymbol{Q}}\|$ (the Euclidean norm or length of the vector) whereas the angle (**E,H,M**) is intended as the trigonometric angle (in the interval $[-\pi, +\pi]$) formed by each vector $\vec{\boldsymbol{T}}_{\boldsymbol{Q_i}}$ with the unit directions $\boldsymbol{e}_i$ (see subpanels A-B). **C,F,I)** Time-evolution (Mean±S.E.M.) of $\|\vec{\boldsymbol{T}}_{\boldsymbol{Q}}\|$ for non-transformed epithelia in the whole (**C**), inner (**F**) and outer (**I**) elliptical quadrants. **D,G,L)** Time-evolution (Mean±S.E.M.) of $\|\vec{\boldsymbol{T}}_{\boldsymbol{Q}}\|$ for *KRAS*-transformed epithelia in the whole (**D**), inner (**G**) and outer (**L**) elliptical quadrants. **E,H,M)** Time-evolution (Mean±S.E.M.) of the angle between vector $\vec{\boldsymbol{T}}_{\boldsymbol{Q_i}}$ with the unit directions $\boldsymbol{e}_i$ in each elliptical quadrant $Q_i$ for non-transformed and *KRAS*-transformed epithelia in the whole (**E**), inner (**H**) and outer (**M**) elliptical quadrants. **N-O)** Time-evolution (Mean±S.E.M.) of force-resultant magnitudes from opposite elliptical quadrants (see subpanels A-B) – i.e. $\|\vec{\boldsymbol{T}}_{\boldsymbol{Q_1}} + \vec{\boldsymbol{T}}_{\boldsymbol{Q_3}}\|$ **and** $\|\vec{\boldsymbol{T}}_{\boldsymbol{Q_2}} + \vec{\boldsymbol{T}}_{\boldsymbol{Q_4}}\|$ (also simply referred to as $\vec{\boldsymbol{T}}_{DIPOLES}$) –is plotted in colour code, according to the legend: **T)** for non-transformed circular epithelia; and, **U)** for *KRAS*-transformed circular epithelia. Statistics over 15 non-transformed epithelia from at least 4 independent experiment repeats and 3 KRAS-transformed epithelia from two different repeats. Median over each epithelial domain at each time point of its evolution.

**Fig. S8.**

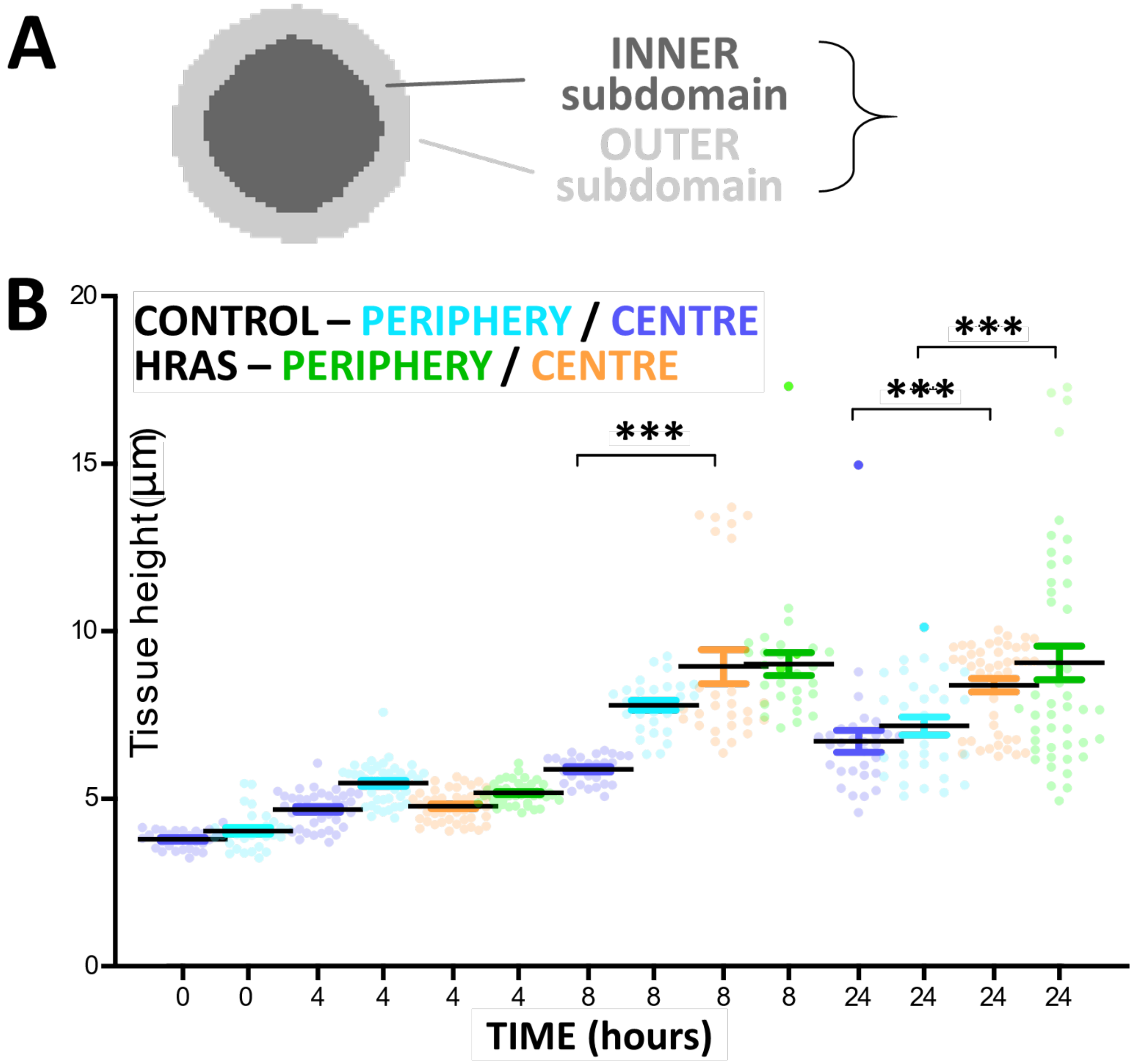


**Epithelial thickness in the inner and outer epithelial domains.**
**A)** Schematic representing the whole epithelial domain along with its outer and inner subdomains. **B)** MEAN±S.E.M. of measurements from at least 3 individual patterns. Kruskal-Wallis statistic test with Dunn's Multiple Comparison Test ***p<0.001.

**Fig. S9.**

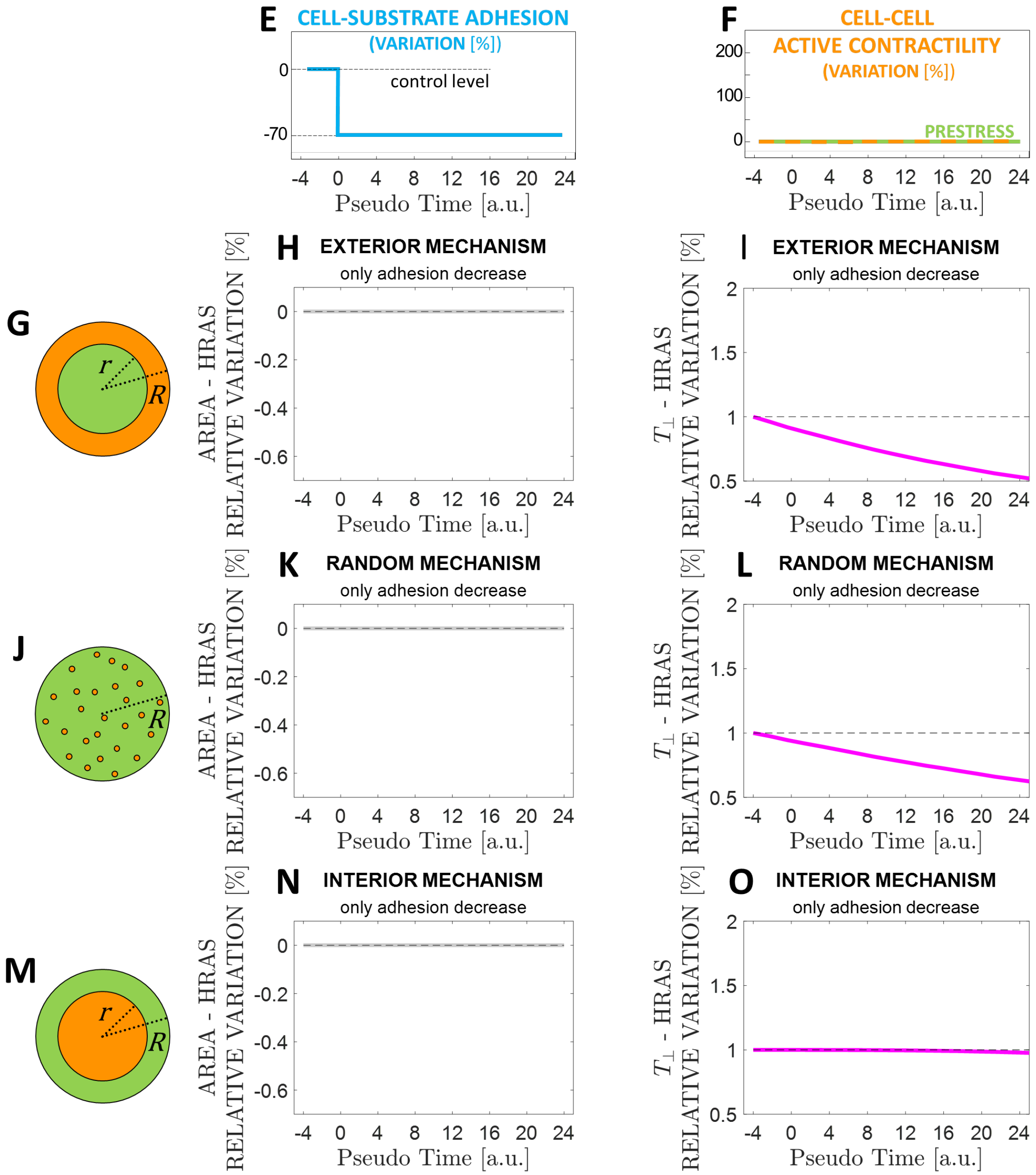


**In silico model predictions in the limit case of sole variation of cell-substrate adhesion (no variation in cell-cell active contractility). A-D)** See Figure 6 A-D for a schematic of the model. **E)** Adhesion with substrate stays at pre-transformation levels in correspondence with the epithelial subdomain $\Omega_1$ that is subject to pre-strain only whereas it decreases by 70% in correspondence with the epithelial domain $\Omega_2$, which is subject to additional active tension. **F)** Active intercellular tension does not increase with respect to pre-transformation levels in the epithelial subdomain $\Omega_2$ and, thus, it stays constant in the whole epithelial domain. **G,J,M)** Schematics illustrating the different topologies according to which active tension can locally increase within the tissue (subdomains $\Omega_1$ and $\Omega_2$ are color-coded in green and orange respectively). **H,K,N)** Epithelial surface area trends in correspondence to each of the scenarios in panels G-I respectively. **I,L,O)** Epithelial traction force trends in correspondence to each of the scenarios in panels G-I respectively.

**Fig. S10.**

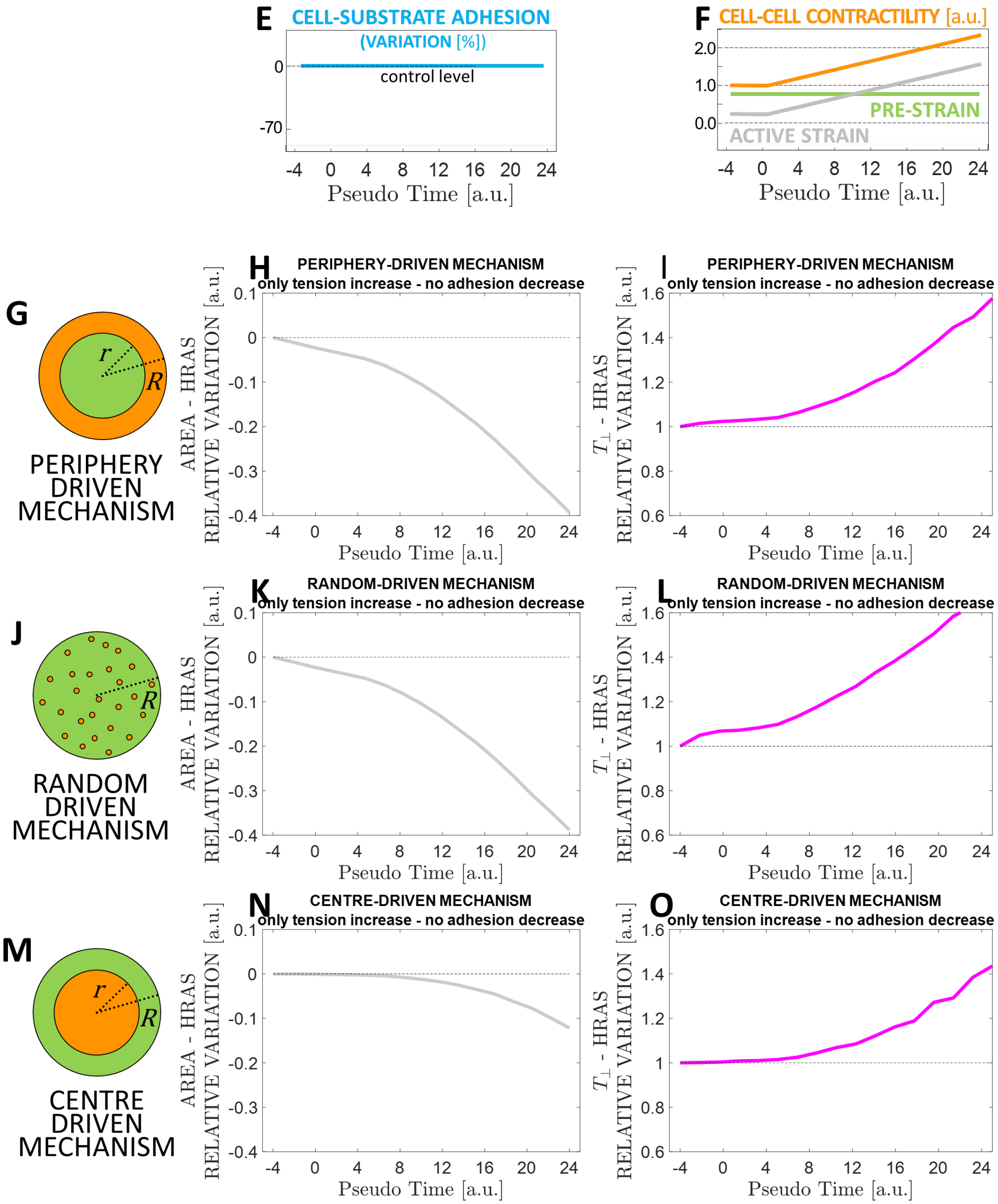


**In silico model predictions in the limit case of sole variation of cell-cell active contractility (no variation in cell-substrate adhesion). A-D)** See Figure 6 A-D for a schematic of the model. **E)** Adhesion with substrate stays at pre-transformation levels in correspondence of the whole epithelial. **F)** Active intercellular tension increases monotonically within the epithelial subdomain $\Omega_2$. **G,J,M)** Schematics illustrating the different topologies according to which active tension can locally increase within the tissue (subdomains $\Omega_1$ and $\Omega_2$ are color-coded in green and orange respectively). **H,K,N)** Epithelial surface area trends in correspondence to each of the scenarios in panels G-I respectively. **I,L,O)** Epithelial traction force trends in correspondence to each of the scenarios in panels G-I respectively.

**Table S1.**

| | **$t$=-4h** | **$t$=0** | **$t$=24h** |
|---|---|---|---|
| **cell-matrix adhesion weakening factor $\alpha$** | 1.0 | 0.3 | 0.3 |
| **epithelial baseline contractile pre-strain $\varepsilon_0^c$** | -0.5 | -0.5 | -0.5 |
| **cell active contractile strain $\varepsilon^c$** | -0.15 | -0.15 | -1.0 |
| **total cell-cell contractile strain $\varepsilon = \varepsilon_0^c + \varepsilon^c$** | -0.65 | -0.65 | -1.50 |

In silico adhesion and contractility parameters.

**Movie S1.**

**Time-lapse of phase-contrast images of non-transformed MCF10A epithelial monolayer.**
Micropatterned MCF10A ER:HRAS treated with DMSO (control) monitored over 50 hours (scale bar = 100 μm).

**Movie S2.**

**Time-lapse of phase-contrast images of HRAS-transformed MCF10A epithelial monolayer.**
Micropatterned MCF10A ER:HRAS treated with 4-OHT (RAS-transformed) monitored over 50 hours (scale bar = 100 μm).

**Movie S3.**

**Time-lapse of traction maps of non-transformed MCF10A epithelial monolayer.**
Colour map represents the component normal to the centre of the monolayer, imposed on phase contrast images (scale bar = 100 μm). Total duration: 50 hours.

**Movie S4.**

**Time-lapse of traction maps of HRAS-transformed MCF10A epithelial monolayer.**
Colour map represents the component normal to the centre of the monolayer, imposed on phase contrast images (scale bar = 100 μm). Total duration: 50 hours.
